# Comparing hormone dynamics in cereal crops via transient expression of hormone sensors

**DOI:** 10.1101/2023.11.14.567063

**Authors:** Thai Q. Dao, Colleen Drapek, Alexander Jones, Samuel Leiboff

## Abstract

Plant hormones are small molecules which elicit profound physiological responses. Although plant hormone biosynthesis and response genes have been critical for agricultural improvement, it has been difficult to experimentally compare hormone biology across species because of complex phenotypic outputs. We used transient expression of genetic hormone sensors and transcriptomics to quantify tissue-specific gibberellic acid (GA) and auxin responses across tissues and genotypes in cereal crops. We found that the FRET-based GPS2 biosensor detects exogenous GA treatments in maize, barley, sorghum, and wheat, in both vegetative and floral tissues. Measuring GPS2 output across GA dosages revealed tissue- and genotype-specific differences in GA sensor response. We observed marked differences in maize vs barley leaves and floral tissues and an unexpected drop in GPS2 output in the maize *d1* GA biosynthesis mutant after GA treatment, likely reflecting differences in bioactive GA content, GA transport, and mechanisms of GA response. We then used RNAseq to measure transcriptional responses to GA treatment in leaves from maize wildtype, *d1*, and barley as well as floral tissues from maize and barley for a cross-tissue, cross-genotype, and cross-species GA-response comparison. After orthology prediction and analysis of within- and cross-species GO-term enrichment, we identified core sets of GA-responsive genes in each species as well as maize- barley orthogroups. Our analysis suggests that downregulation of *GA-INSENSITIVE DWARF1* (*GID1*) and upregulation of α*-Expansin1* (*EXPA1*) orthologs comprises a universal GA-response mechanism that is independent of GA biosynthesis, and identifies F-Box proteins, hexokinase, and AMPK/SNF1 protein kinase orthologs as unexpected cross-tissue, cross-genotype, and cross-species GA-responsive genes. We then compared the transient expression of the DR5, DR5v2, and DII-mDII auxin reporters in barley and maize and find that although DR5 did not respond to exogenous auxin in barley, DR5v2 responded to auxin treatment with a similar magnitude as in maize. Both species display auxin-mediated DII degradation that requires the 26S proteasome.

## INTRODUCTION

Cereal crops form the bedrock of our global agricultural system. The Green Revolution spurred a substantial increase in yield and increased food security for millions of people, based on the adoption of high yielding varieties (HYVs) of food crops (Gollin et al. 2021). Many HYVs exhibit reduced stature and increased grain production, which we now know are consequences of genetic alterations in the gibberellin biosynthesis and signaling pathway (Fu et al. 2002, Peng et al. 1999, Sasaki et al., 2002, Ueguchi-Tanaka et al., 2005).

Gibberellins (GAs) are a family of tetracyclic diterpenoid molecules that are involved in a wide range of developmental and physiological processes in plants, including root and shoot elongation, induction of flowering, fruit production, and seed germination (Brian 1959, El-Sharkawy et al. 2017, Fleet and Sun 2005, Serrani et al. 2007, Shani et al. 2012). Of the more than 130 naturally occurring GAs discovered so far, only a few have been found to be bioactive: GA1, GA3, GA4, and GA7 (Yamaguchi 2008).

Due to the integral role that GAs play in plant growth, their metabolism and signaling mechanism have been heavily investigated using the mouse-ear cress *Arabidopsis thaliana* as a model system. Cellular GA concentration is a product of biosynthesis, catabolism, import and export (Sun et al. 1994, Binenbaum et al. 2018). In brief, the GA biosynthetic pathway requires the conversion of geranylgeranyl pyrophosphate (GGPP) to ent-kaurene in the plastid, catalyzed by three genes *GA1*, *GA2*, and *GA3*, which encode an ent-copalyl diphosphate synthase, an ent-kaurene synthase, and an ent-kaurene oxidase, respectively (He et al. 2020, Sun et al. 1994). In the endoplasmic reticulum, ent-kaurenoic acid oxidase (KAO), a member of the cytochrome P450 family, catalyzes the subsequent conversion of ent-kaurene to GA12 and/or GA53 (Helliwell et al. 1999, Helliwell et al. 2001). The conversion of GA12 into bioactive GA species in the cytoplasm involves a multi-step process, catalyzed by several enzymes including GA 20-oxidases (encoded by *GA20ox* genes) and GA 3-oxidases (encoded by *GA3ox* genes) (He et al. 2020, Olszewski et al. 2002). In addition, three classes of enzymes have been identified to catabolize bioactive GAs to maintain homeostasis, including GA 2-oxidases (*GA2ox* genes), GA methyltransferases (*GAMT* genes), and GA 16,17-oxidases (*GA16,17ox* genes) (Rieu et al. 2008, Thomas et al. 1999, Varbanova et al. 2007, Zhang et al. 2011, Zhu et al. 2006). GA is also subject to differential import based on variable membrane permeability amongst GAs as well as active import by NPF3 and AtSWEET13/14 (Tal et al. 2016, Kanno et al. 2016). Although protein-mediated GA efflux is anticipated based on long-distance GA mobility and environmental regulation of efflux rates (Mertz 1985), no GA efflux carrier has been identified (Binenbaum et al. 2018).

Transduction of GA signaling is mediated by the GA receptor GIBBERELLIN INSENSITIVE1 (GID1) and DELLA proteins such as Arabidopsis GIBBERELLIC ACID INSENSITIVE (AtGAI), REPRESSOR OF THE *ga1-3* MUTANT (AtRGA), or maize DWARF8 (D8) and DWARF9 (D9) (Lawit et al., 2010, Peng et al., 1997, Silverstone et al., 1998). In the absence of GA, DELLAs physically interact with transcription factors such as PHYTOCHROME INTERACTING FACTOR3 (PIF3), effectively preventing them from activating transcription. Upon binding to GA, GID1 undergoes a conformational change that promotes its stable interaction with DELLAs, followed by the recruitment of F-box proteins such as GIBBERELLIN INSENSITIVE2 (GID2) as a part of the E3 ubiquitin ligase SCF complex. The resulting ubiquitination and degradation of DELLAs release PIF3 to promote transcription of GA-responsive genes, including modulators of cell elongation and cell division, interaction with other hormone pathways, other transcription regulators, as well as feedback inhibition of GA biosynthesis (Davière and Achard, 2013, Ogawa et al. 2003). Previous transcriptomic analyses showed down-regulation of *GA20ox* and *GA3ox* genes, as well as up-regulation of *GA2ox* genes in response to higher endogenous GA level and GA treatment (Ci et al. 2021, Ogawa et al. 2003, Olszewski et al. 2002, Tan et al. 2018).

Despite extensive research into GA signaling and metabolism, little is known about the distribution of GA within plant tissues, especially cereal crops. Measurements of GA extracts from bulk tissue by gas chromatography-mass spectrometry revealed changes in internal GA content in response to light affect germination and elongation (Hirose et al. 2012, Jiang et al. 2016, Zhao et al. 2007). However, GA dynamics at the cellular and tissue levels, and how GA contributes to tissue patterning and environmental response were previously limited by a lack of biosensors that can be observed in real-time. A green fluorescent protein (GFP) - REPORESSOR OF ga1-3 (RGA) fusion protein that is degraded in the presence of GA showed that DELLA accumulation is influenced by iron content in growth media (Silverstone et al. 2001, Wild et al. 2016). More recently, the GIBBERELLIN PERCEPTION SENSOR1 (GPS1) was developed based on Förster Resonance Energy Transfer (FRET) as a direct biosensor for GA (Rizza et al. 2017). GPS1 is a single fusion protein composed of an AtGAI truncation fused to a yellow fluorescent protein (enhanced dimerization Aphrodite or edAFP) linked with the GA receptor AtGID1C fused to a cyan fluorescent protein (enhanced dimerization Cerulean or edCer). GPS1 is capable of detecting GA gradients in *Arabidopsis* shoot and root tissues, as well as changing GA levels either by exogenous GA application or light conditions (Rizza et al. 2017) or via manipulation of enzyme expression (Rizza et al. 2021). Other ratiometric hormone reporters have also been developed in *Arabidopsis*, including the auxin-sensing DR5v2/DR5 and R2D2 reporters (Liao et al. 2015).

Given the lack of tools to study GA biology in grass models, we aimed to test the function of GPS as a GA biosensor in a variety of cereal grains. In this report, we demonstrate that GPS2, a second-generation GPS with improved orthogonality and reversibility (Griffiths et al. 2023, Drapek et al. 2023), responds to exogenous GA applications in transiently transfected leaf and floral tissues from maize, sorghum, barley, and wheat. Using nuclear localized nlsGPS2, we detected differential GA accumulation between leaf and floral tissues of maize B73 and barley Golden Promise as well as profound differences in GA accumulation between tissues of maize B73 and the GA-deficient *dwarf1* mutant. Pursuing these differences with mRNA-seq in a cross-tissue and cross-species analysis revealed fundamental differences in how the maize B73 and *d1* transcriptomes respond to GA treatment, as well as conserved elements of GA response between maize and barley. Our studies not only validate GPS2 as a functional GA sensor in agricultural models, but also demonstrate transient expression via particle bombardment as an effective tool for comparative studies of hormone biology in grasses in a short timeframe.

## METHODS

### Plant growth conditions

Maize (*Zea mays*) inbred B73 is available through the US National Plant Germplasm system under accession PI 550473. Plants were grown in 2-gallon pots fertilized with 20-20-20 NPK, at the density of 4 plants/pot, in greenhouse conditions with 16 hour LED-supplemented days, controlled temperature ranging from 18-24°C with watering twice a week.

The *dwarf1* maize line used in this study contains the *d1-6016* allele in the B73 inbred background (Chen et al. 2014, GRIN accession MGS 14087), and was a gift of Sarah Hake. Plants were grown in 2-gallon pots fertilized with 20-20-20 NPK, 4 plants/pot, in greenhouse conditions with 16 hour LED-supplemented days, controlled temperature ranging from 18-24°C with watering twice a week.

Barley (*Hordeum vulgare*) cultivar Golden Promise is available through the US National Plant Germplasm system under accession PI 343079. Plants were grown in 48-well starter trays fertilized with 20-20-20 NPK, 1 plant/well, in greenhouse conditions with 16-hour LED-supplemented days, controlled temperature ranging from 18-24°C with daily watering.

Wheat (*Triticum turgitum*) cultivar Kronos is available through the US National Plant Germplasm system under accession PI 576168. Plants were grown in 48-well starter trays fertilized with 20- 20-20 NPK, 1 plant/well, in greenhouse conditions with 16-hour LED-supplemented days, controlled temperature ranging from 18-24°C with daily watering.

Sorghum (*Sorghum bicolor*) inbred BTx623 is available through the US National Plant Germplasm system under accession PI 564163. Plants were grown in 2-gallon pots fertilized with 20-20-20 NPK, 4 plants/pot, in greenhouse conditions with 16-hour LED-supplemented days, controlled temperature ranging from 18-24°C with watering twice a week.

### Construct Information

nlsGPS1-NR was previously described (Rizza et al., 2017), The monocot nlsGPS2 sensor is based on the sequence for the next generation nlsGPS2 sensor, which contains four charge exchange mutations at the interface between *At*GID1C (K13E and K28E) and *At*GAI (E51K and E54R), a change that improves orthogonality by reducing interaction with endogenous GID1 and DELLA proteins whilst maintaining intermolecular interactions in the biosensor. Experiments in vitro show a similar ratio change in response to GA4 and improved reversibility (BioRxiv version pending).

The amino acid sequence for the monocot nlsGPS2 was optimized for expression in barley and synthesized (Genewiz). The nls sequence, donor, acceptor, AtGID1C, linker, and AtDELLA fragments were codon optimized using GeneArt (ThermoFisher) including BpiI overhangs for assembling by GoldenGate (Engler et al 2014). The fragment was designed with additional sequence adapters for S site overhangs (5’) and C site overhangs (3’) for assembling via GoldenGate cloning using the MoClo kit described in Engler et al. 2014. The final assemblage contained an S overhang (5’) and C overhang (3’) as well as for assembling into a level 1 position with the same fragments described above and amino acid scar sequences between some fragments from GoldenGate assembling (LE between nls and acceptor Aphrodite, GG between the linker and AtDELLA-based fragment, GG between the linker and AtGID1C-based fragment). A level 1 sequence was assembled with the *Zea mays* UBIQUITIN1 promoter sequence (983 bp) and first UBIQUITIN1 intron driving sensor expression (level 0 promoter sequence) and the 35S terminator sequence (level 0 terminator sequence). A final binary destination vector level 2 construct was assembled using a Hygromycin selection cassette (position1), an insulator sequence (position2) and the monocot codon-optimized ZmUBI::nlsGPS2::t35S in position 3 using level 2 acceptor pAGM4723.

The monocot nlsGPS-NR control sensor amino acid sequence contains the same four charge exchange mutations as the nlsGPS2 but with the additional NR mutations (S114A and F115A in the AtGID1C binding pocket). DR5v2-ntdTomato/DR5-n3GFP and R2D2 were previously described (Liao et al. 2015) and obtained from Addgene (DR5v2/DR5 catalog #61628, R2D2 catalog #61629).

### Biolistic Bombardment and Hormone Treatment

For bombardment of leaves, tissues were collected at the following conditions: up to 8 youngest leaves from maize plants at 60 days after germination (DAG); up to 4 youngest leaves from barley at 40-60DAG; up to 4 youngest leaves from wheat at 40-60DAG; up to 8 youngest leaves from barley at 60DAG. Tissues from up to 3cm from the base of the SAM were collected and dissected into approximately 5mm x 5mm pieces.

For bombardment of floral tissues, florets were collected from the tassel of maize plants at 90DAG, and main inflorescence of barley plants at 50DAG, at which point silk organs were clearly visible. Individual florets from up to 3cm from the base of the inflorescence were collected.

Plants were dissected using a scalpel with the aid of a Leica DMi8 microscope at 10x magnification. Tissues were arranged in concentric circles on petri dishes containing Murashige and Skoog (MS) growing medium supplemented with 0.4M D-sorbitol, 1% agar, pH 5.7. The centers of the circles were left empty up to a radius of 2cm to minimize tissue damage from bombardment. Tissues were bombarded within 4 hours of dissection.

Microprojectile bombardments were performed as described by Kirienko et al. 2012. DNA was prepared by commercially available column purification in midiprep format (ZymoPure). DNA concentrations used were between 0.5-2ug/uL, with no attempt to maximize transfection efficiency based on DNA concentration.

All bombardments were performed with Bio-Rad Tungsten M-10 microcarrier, prepared as follows. 20 mg of microcarriers were weighed in a 1.5 mL centrifuge tube and washed by mixing in HPLC grade 70% ethanol for 5 minutes, soaked for 15 minutes, and then pelleted by microcentrifugation for 10s. Pellets were washed by mixing in 1mL of sterile distilled water for 1 minute, centrifuged and liquid was discarded for a total of three times. Finally, microcarriers were resuspended in sterile 50% glycerol, divided into 50uL aliquots, and stored at -20°C until use.

Each 50uL aliquot of microcarriers was vortexed for 5 minutes and the following were added sequentially: 20uL of 0.1M spermidine with mixing 1 minute; 6 uL of DNA at 0.5-2 ug/uL with mixing for 1 minute; and 50uL of 2.5M CaCl2 with mixing for 5 minutes. Microcarriers and DNA were centrifuged for 15s and supernatant was discarded. Pellets were washed once with 140 uL of 70% ethanol, and then again with 140 uL of 100% ethanol. Microcarriers were then resuspended in 50 uL of 100% ethanol. For each bombardment, 8 uL of microcarriers in 100% ethanol was pipetted onto a macrocarrier in a sterile hood and left to evaporate.

All bombardments were performed with a PDS-1000 (Bio-Rad, CA) system with the following settings: 1100 psi rupture disk (Bio-Rad, CA); 6 cm stopping distance between the plate and the macrocarrier stopping screen; and vacuum pressure of at least 25 inHg prior to firing. Helium pressure at the tank was regulated at 1600 psi.

After bombardment, tissues were treated with hormones as follows. For GA treatment, 1 mL of filter-sterilized 0.4M D-sorbitol in water supplemented with 0, 0.1, 1, 10, or 100uM GA3 were pipetted onto tissue samples. Tissue was allowed to recover for 16 to 24h at room temperature in the dark prior to imaging. For IAA treatment, tissue was first allowed to recover for 16 to 24h at room temperature in the dark, followed by adding 1mL of sterile water or 100uM of MG-132 in sterile water and incubated for 1h in the dark. Afterwards, any remaining water or MG-132 were discarded, followed by adding 1mL of sterile water or a combination of 0uM IAA and 100uM MG-132, 10uM IAA and 0uM MG-132, or 10uM IAA and 100uM MG-132 and incubated for 3h in the dark prior to imaging.

### Image collection

Epifluorescence imaging was performed on a Leica DMi-8 using a HC PL APO 20x dry 0.80 NA objective. For GPS2 and GPS1-NR, images were collected using a CYR 71010 quad band filter with the following optical bands: Excitation: 422-450; 495-517; 566-590; 710-750, Dichroic: 459; 523; 598; 763, Emission: 462-484; 527-551; 602-680; 770-850. CFP was excited using a 440 nm LED and specifically collected using a 460/80 emission filter wheel and the CYR 71010 quad band filter cube; YFP was excited using a 510 nm LED and specifically collected using a 535/70 emission filter wheel and the CYR 71010 quad band filter cube; FRET was excited using a 440 nm LED and specifically collected using a 535/70 emission filter wheel and the CYR 71010 quad band filter cube. For DR5v2/DR5, images were collected with two channels: ntdTomato was excited using a 575 nm LED and specifically collected using a 590/50 emission filter wheel and the CYR 71010 quad band filter cube; GFP was excited using a 470 nm LED and collected at 527/15 using a FITC filter cube. For R2D2, images were collected with two channels: ntdTomato was excited using a 575 nm LED and specifically collected using a 590/50 emission filter wheel and the CYR 71010 quad band filter cube; YFP was excited using a 510 nm LED and specifically collected using a 535/70 emission filter wheel and the CYR 71010 quad band filter cube. The z-stacks were acquired with a step-size of 5 um.

### Image analysis

Nuclei segmentation and measurement of GPS2 emission ratio were performed in ImageJ (Fiji). A macro was written to automatically process each z-stack following the protocol described by Rizza et al. 2019 with a few modifications. Z project was first performed on each stack, with “Sum slices” used as the option, followed by Subtract background (rolling ball radius = 25 pixels) and Gaussian blur (sigma = 2). The stack was split into individual channels for CFP, FRET, and YFP. The FRET and YFP channel were duplicated, converted to 8-bit format, binarized using Auto-Local Threshold (method = Bernsen, radius = 25, parameter 1 = 0, parameter 2 = 0, white stack), and then multiplied together using Image Calculator to create a binary mask containing the nuclear segments. Subsequently, the original FRET channel was divided over the CFP channel using Image Calculator to create a new 32-bit image containing the emission ratio values. This emission ratio image was multiplied with the previously generated mask and divided by 255. For selection of nuclear regions of interest (ROIs), the emission ratio was duplicated and binarized by thresholding (min = 0.1) and pixel erosion (radius = 3). A selection was created over the entire binary mask and split into individual ROIs in the ROI Manager panel. All ROIs were multi-measured on the initial emission ratio image with the following parameters: area, mean gray value, standard deviation, and shape descriptors (circularity, aspect ratio, roundness, solidity).

Nuclear segmentation of DR5v2/DR5 and R2D2 was performed as described above with the following modifications. The stack was split into two individual channels, both of which was duplicated, converted to 8-bit format, binarized, and multiplied together to create a binary mask containing the nuclear segments. A pixel erosion (radius = 3) was applied to the mask, which was then selected and split into individual ROIs in the ROI Manager panel. All ROIs were multi- measured on each of the individual channels with the following parameters: area, mean gray value, standard deviation, and shape descriptors (circularity, aspect ratio, roundness, solidity). For R2D2, the mean gray values of the ntdTomato channel were subsequently divided over those of the YFP channel to obtain the mDII/DII emission ratio.

Subsequent statistical analyses were performed in RStudio using R version 4.1.3. ROI measurements were further filtered by shape (circularity > 0.4) and GPS ratio (mean gray value < 10) to remove outliers. We calculated count, mean, standard deviation, standard error, and 95% confidence interval using the summarySE() function in the package Rmisc v1.5. We tested the relationship between increasing dosages of GA_3_ treatment and GPS emission ratio by ANOVA using the aov() function, and Tukey’s Honest Significance Difference test using the TukeyHSD() function in the package stats v4.1.3. We calculated the effect size of each treatment to GPS emission ratio by Cliff’s Delta using the cliff.delta() function in the package effsize v0.8.1.

### Tissue collection for RNAseq

For bulk RNA-seq library construction, leaf tissues were collected at the following conditions: up to 8 youngest leaves from maize B73 and d1 plants at 54DAG; up to 8 youngest leaves from maize B73 plants at 90DAG; up to 4 youngest leaves from barley at 45DAG. Leaf tissues from up to 3cm from the base of the SAM were collected. Floret tissues were collected from the tassel of maize B73 plants at 90DAG and main inflorescence of barley at 45DAG, from up to 3cm from the base of the inflorescence. Tissues were dissected into approximately 5mm x 5mm pieces were dissected using a scalpel with the aid of a Leica M205 FCA microscope at 10x magnification.

For GA treatment, tissues were arranged on petri dishes containing Murashige and Skoog (MS) growing medium supplemented with 0.4M D-sorbitol, 1% agar, pH 5.7. 1 mL of filter-sterilized 0.4M D-sorbitol in water supplemented with 0 or 100uM GA3 were pipetted onto tissue samples. Tissue was allowed to incubate for 16 at room temperature. Tissues were then collected and sealed in 1.5mL eppendorf tubes containing 3 stainless steel beads, flash frozen in liquid nitrogen, and stored at -80C until extraction.

### RNA extraction and quantification

RNA extraction was performed as previously described (Leiboff and Hake, 2019). Frozen tissues were homogenized using a SPEX Geno Grinder 2010 automated tissue homogenizer (freq = 25 Hz, duration = 30-45s), then returned to liquid nitrogen. RNA was extracted using TRIzol, precipitated using 2-propanol, washed with 70% ethanol, and resuspended in nuclease- free water. RNA was quantified using a QuBit BR-RNA kit and potential RNA degradation evaluated on a 1% agarose gel in 1x TAE buffer. Total RNA concentration was determined using a Qubit Flex Fluorometer and Qubit RNA BR reagents (Thermo Fisher Scientific).

### Library preparation and quantification

Approximately 1 ug of RNA was used as input to the NEBNext Poly(A) mRNA magnetic isolation module (New England Biolabs, catalog #E7490S), followed by NEB Ultra II RNA library prep kit for Illumina (New England Biolabs, catalog #E7770S, E7335S, E7500S, E7710S).

Multiplexed library synthesis was carried out according to manufacturer specifications with 8 cycles of PCR enrichment as recommended by the manufacturer. Libraries were purified using SPRIselect beads (Beckman Coulter, catalog #B23318). Library quality and quantity was verified by Agilent 2100 BioAnalyzer Instrument, and qPCR by the Oregon State University Center for Quantitative Life Sciences (CQLS).

### RNA sequencing, alignment, and normalization

All libraries were added in an equimolar mix and sequenced using one lane S4 Illumina NovaSeq 6000 with 150 bp paired-end sequencing chemistry at University of Oregon, Genomics and Cell Characterization Core Facility (GC3F). Sequence quality control and read processing were performed with fastp v0.20.1. Reads were aligned to the maize B73- REFERENCE-NAM-5.0 genome or the barley MorexV3 pseudomolecule assembly (minimum intron length 60, maximum intron length 50000) using HiSAT2 v2.2.1. Aligned reads were counted using the featureCounts_unionExon software in the package Subread v2.0.1 to the maize B73-REFERENCE-NAM-5.0 gene set or the barley MorexV3 gene set. Differential gene expression analysis from raw counts was performed with R package DESeq2 v1.34.0. Genes with less 5 counts (or 1 read per million) in less than 6 libraries for maize, or less than 3 libraries for barley were excluded from subsequent analyses.

### Differential gene expression analysis

All statistical analyses were performed in Rstudio using R version 4.1.3 unless otherwise noted. The FPKM (Fragments Per Kilobase of exon model per Million mapped reads) of each gene was calculated based on the number of read counts mapped and the length of the union-exon transcript reported by featureCounts. For comparison between mock and GA3-treated tissues, log2 fold change of each gene was calculated using the results() function in DESeq2 v1.34.0. A gene was considered differentially expressed in response to GA if its log2 fold change was > 0.58 (equivalent to 1.5 fold change), with an adjusted p-value < 0.05. Heatmaps of differentially expressed genes were generated using the pheatmap() function in pheatmap v1.0.12.

### GO-term analysis

For Gene Ontology (GO) enrichment in maize tissues, an organism package for *Zea mays* was made using the makeOrgPackage() function in the package AnnotationForge v1.36.0 (Carlson & Pagès 2022). GO enrichment analyses of DEGs were conducted using the enrichGO() function in the package enrichplot v1.14.2, using the Benjamini-Hochberg procedure with a p-value cutoff of 0.01 and q-value cutoff of 0.05 to control the false discovery rate (Yu 2022).

### Comparison of syntenic orthologs

Orthogroups between maize and barley were identified on the Linux-based OrthoFinder platform (Emms and Kelly, 2019). Protein sequences from the maize B73-REFERENCE-NAM-5.0 and barley MorexV3 genomes were used, as well as the rice IRGSP-1.0 and Arabidopsis TAIR10 genomes as outgroups. For correlational analysis of GA response, a dataset was composed of only orthogroups containing a single gene from each species and their corresponding log2 fold change values. A correlation matrix was generated using the correlate() function in the package corrr v0.4.3.

### Data and code availability

The raw sequence data used in this study are available in the NCBI Sequence Read Archive (SRA) within BioProject PRJNA997436.

## RESULTS

### GPS2 responds to GA treatment in leaf and floral tissues of multiple species

To assess whether GPS is a functional GA sensor in grasses, we transiently transformed the nuclear targeted GPS2 (nlsGPS2), an improved version of nlsGPS1 under the control of the maize ubiquitin (ZmUBI) promoter, into leaf and floral tissues using particle bombardment. At 16-24h post-bombardment, nlsGPS2 signal was observed in all tissues tested across all three collection channels (CFP, FRET, and YFP), and could be differentiated from background noise and bombardment artifacts, which were only detected in the CFP and FRET channels (Fig. 1A). The construct accumulated in spherical puncta resembling nuclei in almost all transfected cells, with a few exhibiting both nuclear and membrane fluorescence indicating possible nuclear membrane damage or cell death (Fig. 1A). Data were collected from all nuclei within each field of view. Because we used biolistic bombardment to introduce our constructs, we anticipate that most transformed nuclei originate from epidermal or near-epidermal cell layers in all samples.

**Figure 1.**
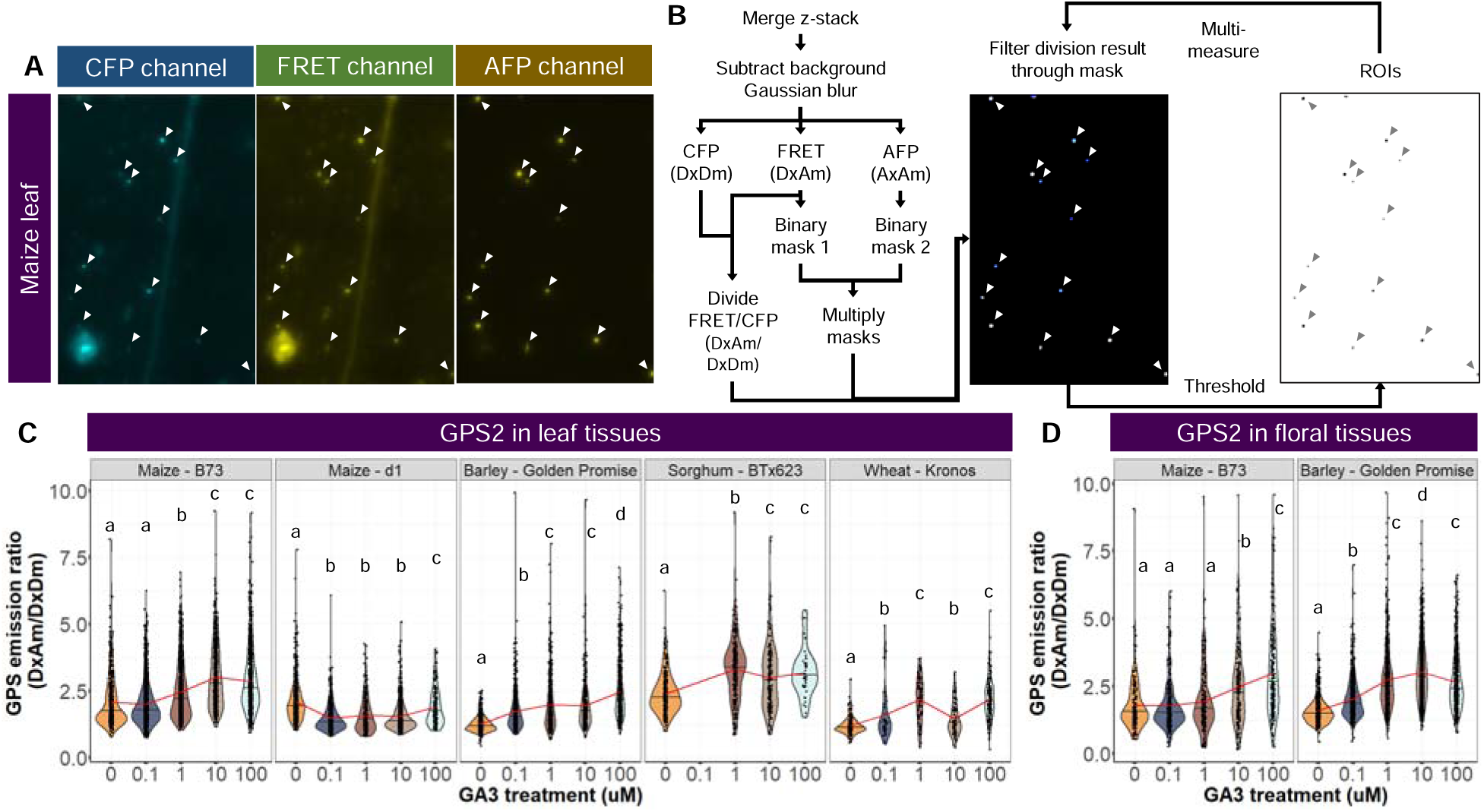
GPS2 transient expression reveals genotype and tissue differences in FRET ratio after GA treatment. A, Transient expression of GPS2 after biolistic bombardment yields characteristic nuclear signal of CFP (left), CFP to AFP FRET ex/em (middle), and AFP ex/em (right). B, Fluorescence quantification based on mask multiplication between binarized channels allows identification of GPS2-expressing regions of interests (ROIs) and calculation of FRET/CFP emission ratio. C, Leaf tissues transiently expressing GPS2 in maize, barley, sorghum, and wheat display a FRET/CFP emission ratio response when treated with increasing levels of GA. Note that maize *d1*, GA biosynthesis mutant, displays a decrease in FRET ratio. Letters, ANOVA posthoc statistical groups, FRET ratio ∼ GA treatment, p<0.05. D, Floral tissues transiently expressing GFP2 in maize and barley respond to GA, but maize only responds at high GA treatments (10 uM and 100 uM). Letters, ANOVA posthoc statistical groups, FRET ratio ∼ GA treatment, p<0.05. White/grey arrows indicate nuclei and resulting ROIs.

Immediately after bombardment, each tissue plate was incubated with an exogenous GA3 application of various concentrations (0, 0.1, 1, 10 or 100uM) to assess potential GPS2 response to changing GA levels. GPS emission ratio was measured as the fluorescence intensity of FRET divided by that of CFP from each transfected nucleus and averaged over all nuclei on the same image as detailed in the Methods section (Fig. 1B). GPS2 emission ratio was highly variable, as each nucleus underwent an independent transfection event, and thus potentially expressed the construct at different levels relative to background autofluorescence (Fig. 1C,D). Regardless, over thousands of independent observations GPS2 showed an increased fluorescence emission ratio in response to increasing dosages of GA3 application in leaf tissues of maize B73, barley Golden Promise (GP), sorghum BTx623, and wheat Kronos, as well as in floral tissues of maize B73 and barley Golden Promise (Table 1). GPS2 response to exogenous GA3 treatment was non-linear and we observed maximal GPS emission ratio in some tissue and treatment combinations, including maize B73 leaf + 10uM GA3, sorghum BTx623 leaf + 1uM GA3 and barley GP + 10uM GA3, which may represent saturation of the GPS2 sensor based on different starting GA within the tissue or differences in rates of GA import, export, and/or catabolism across species. Interestingly, GPS2 emission ratio in maize *d1* leaf tissues did not increase, but decreased with all GA3 treatments, suggesting that nuclear GA concentration in d1 mutant leaves was lowered by exogenous GA application or specifically lowered within the cells measured in our experiment (Fig. 1B). Patterns of GPS2 response were consistent across two independent replicate batches of plants (Fig. S1A,B, Table S1, S3).

**Table 1.** GPS2 in leaf and floret tissues. . N nuclei observed, mean emission ratio, & Cliff’s delta of 2 combined biological replicates.

| GPS2 in leaf tissues |  |  |  |  |  |  |  |  |  |  |
| --- | --- | --- | --- | --- | --- | --- | --- | --- | --- | --- |
|  | Maize – B73 |  |  |  |  | Maize – *d1* |  |  |  |  |
| [GA<sub>3</sub>] (uM) | 0 | 0.1 | 1 | 10 | 100 | 0 | 0.1 | 1 | 10 | 100 |
| Mean<br>±95%CI | 2.08<br>±0.08 | 2.01<br>±0.05 | 2.47<br>±0.06 | 3.01<br>±0.09 | 2.87<br>±0.07 | 2.11<br>±0.06 | 1.48<br>±0.04 | 1.59<br>±0.08 | 1.54<br>±0.04 | 1.89<br>±0.11 |
| Cliff’s Delta | - | 0.00 | 0.26 | 0.53 | 0.46 | - | -0.59 | -0.47 | -0.55 | -0.21 |
| N | 573 | 1181 | 1055 | 678 | 971 | 674 | 715 | 249 | 761 | 166 |
|  | Barley - Golden Promise |  |  |  |  | Sorghum – BTx623 |  |  |  |  |
| [GA<sub>3</sub>] (uM) | 0 | 0.1 | 1 | 10 | 100 | 0 | 0.1 | 1 | 10 | 100 |
| Mean<br>±95%CI | 1.24<br>±0.02 | 1.74<br>±0.06 | 2.00<br>±0.09 | 1.95<br>±0.09 | 2.47<br>±0.08 | 2.40<br>±0.08 | - | 3.32<br>±0.13 | 3.00<br>±0.16 | 3.19<br>±0.29 |
| Cliff’s Delta | - | 0.53 | 0.57 | 0.67 | 0.88 | - | - | 0.53 | 0.31 | 0.53 |
| N | 666 | 551 | 458 | 365 | 614 | 307 | - | 276 | 210 | 41 |
|  | Wheat - Kronos |  |  |  |  |  |  |  |  |  |
| [GA<sub>3</sub>] (uM) | 0 | 0.1 | 1 | 10 | 100 |  |  |  |  |  |
| Mean<br>±95%CI | 1.19<br>±0.05 | 1.61<br>±0.31 | 2.17<br>±0.18 | 1.45<br>±0.13 | 2.20<br>±0.11 |  |  |  |  |  |
| Cliff’s Delta | - | 0.24 | 0.70 | 0.24 | 0.84 |  |  |  |  |  |
| N | 187 | 42 | 81 | 82 | 172 |  |  |  |  |  |
| GPS2 in floral tissues |  |  |  |  |  |  |  |  |  |  |
|  | Maize – B73 |  |  |  |  | Barley – Golden Promise |  |  |  |  |
| [GA<sub>3</sub>] (uM) | 0 | 0.1 | 1 | 10 | 100 | 0 | 0.1 | 1 | 10 | 100 |
| Mean<br>±95%CI | 1.80<br>±0.13 | 1.79<br>±0.11 | 1.95<br>±0.17 | 2.48<br>±0.20 | 3.00<br>±0.21 | 1.60<br>±0.04 | 2.04<br>±0.05 | 2.75<br>±0.09 | 3.02<br>±0.06 | 2.65<br>±0.08 |
| Cliff’s Delta | - | -0.03 | 0.05 | 0.32 | 0.52 | - | 0.42 | 0.73 | 0.82 | 0.70 |
| N | 188 | 272 | 192 | 196 | 223 | 667 | 693 | 659 | 1124 | 618 |

We also bombarded the GA-insensitive nlsGPS2-NR construct into the same tissue types undergoing the same GA3 treatment as a negative control. nlsGPS2-NR exhibited some response to exogenous GA application in maize B73, maize d1, and barley GP leaf tissues as well as barley floral tissues (Figure S1C,D, Table S2, S4), indicating a fraction of the response of nlsGPS2 might be due to effects other than direct GA sensing, but the response effect size was significantly smaller compared to GPS2 samples in all treatments (Fig. S1, Table S1-4).

Taken together, our bombardment experiments indicate that GPS2 is a functional GA biosensor in a variety of cereal grain models and tissue types, and that GPS2 reveals subtle differences in GA accumulation between species/tissues and a differential GA regulatory mechanism in *d1* maize leaf vs. wild-type.

### RNA-seq analysis between B73 and d1 leaves at 54DAG

To explore the gene regulation responses to GA treatment among different tissue types, we performed a transcriptomic analysis of six sample types and two treatments in triplicate: maize B73 and *d1* vegetative leaves, maize B73 and barley GP at the reproductive stage leaves and floral tissues, all treated with either 0uM or 100uM of GA3. Each sample and treatment combination yielded 18-42 Mb of raw RNAseq reads per replicate with 2 x 150 bp read chemistry (Supplementary File 1). At least 95% of the raw read pairs from each library passed quality and length filtering. Of the clean read pairs, 82-94% were uniquely mapped to either the maize or barley reference genome. Genes were considered expressed if at least five read pairs were mapped to the reference genome in more than 25% of the replicates (six replicates for maize, three replicates for barley). In total we detected the expression of 24,948 out of 39,756 maize gene models and 24,566 out of 81,687 barley gene models.

Hierarchical clustering of normalized FPKM values showed high similarity between replicates of each tissue type and hormone treatment combination with two exceptions: maize B73 90DAG florets ± 100uM GA3, and barley GP 45DAG leaf ± 100uM GA3 (Fig. S2, S3). In each of these exceptions, GA-treated and mock-treated tissues showed a mixed clustering pattern, suggesting that GA3 treatment only affected a small portion of their transcriptomes. Interestingly, age of plants had a stronger effect on the transcriptome than genotype or tissue type in maize, as we observed clustering of B73 54DAG leaf with *d1* 54DAG leaf, and B73 90DAG leaf with B73 90DAG floret.

The maize D1 locus encodes ZmGA3ox2, an enzyme that catalyzes the final step of bioactive GA production (GA9 to GA4, GA20 to GA1, GA20 to GA3, and GA5 to GA3) (Chen et al. 2014). Loss of function *d1* mutants show increased accumulation of GA20 and GA29, and decreased levels of GA1, resulting in a dwarf phenotype that could be rescued by treatments of GA1, GA3, or GA5 at the early seedling stage (Spray et al. 1996). To explore potential transcriptomic responses underlying differences in GA homeostasis regulation between B73 and *d1* leaf tissues as revealed by GPS2, we identified differentially expressed genes (DEGs) in mock versus GA3-treated leaf sections from B73 and *d1* plants at 54DAG. Genes were considered differentially expressed if their |log2 fold change| ≥ 0.58, which corresponds to |fold change| ≥ 1.5, and *q*-value ≤ 0.05. We identified 2,103 total DEGs in response to GA3 treatment in B73 or *d1* leaf tissues. Of these DEGs, 843 (40.1%) were unique to B73 leaf GA treatment, 890 (42.3%) were unique to *d1* leaf GA treatment, and 370 (17.6%) were shared by both during GA treatment (Fig. 2A).

**Figure 2.**
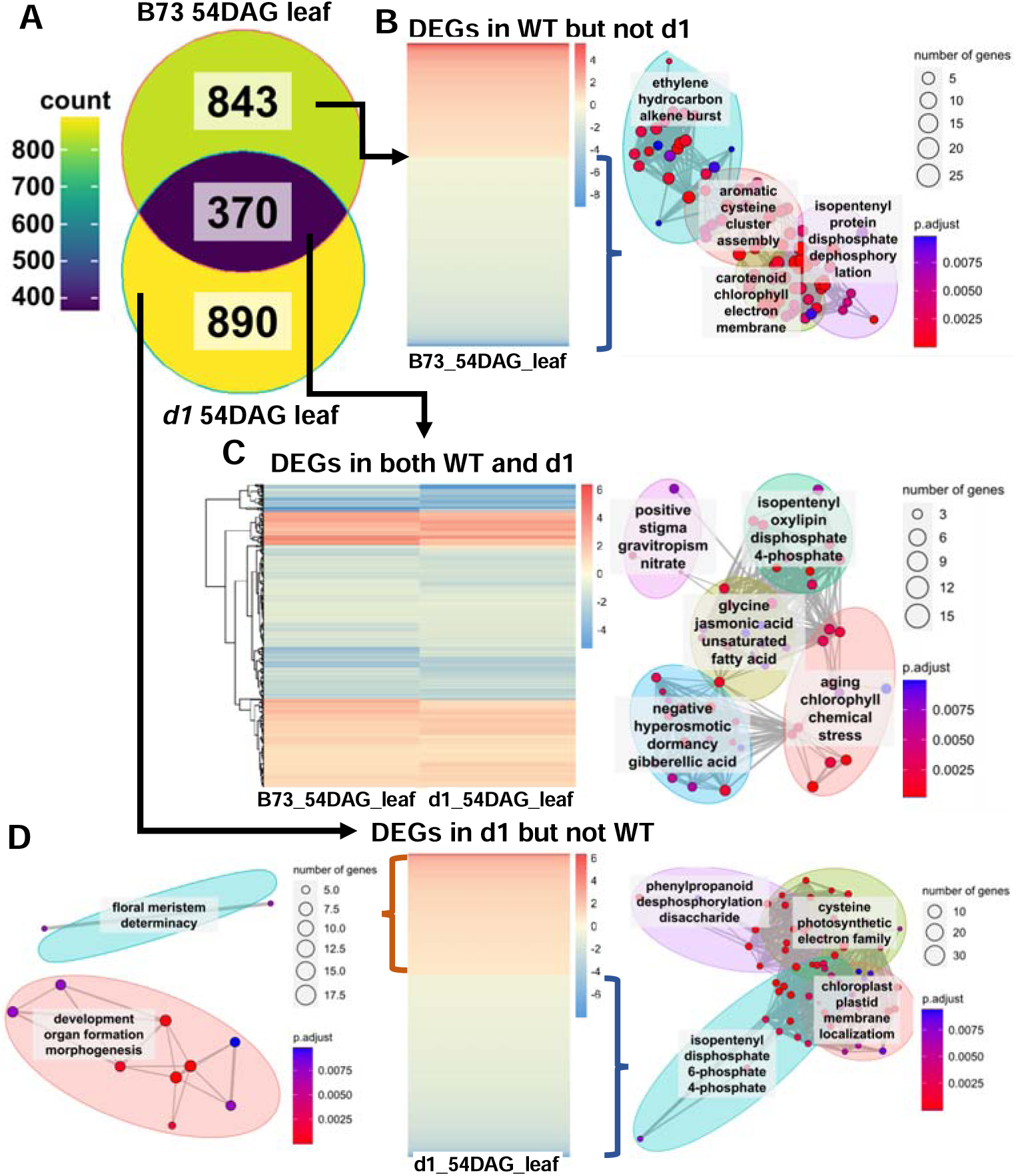
RNAseq of maize B73 and *d1* mutant leaf identifies mostly non-overlapping genes between genotypes in response to GA treatment. A, Venn diagram of DEGs called between B73 and *d1* 54DAG leaf GA treatments identifies a high number of DEGs unique to B73 or *d1* treatments. B, Upregulated genes unique to B73 were not enriched for GO terms, but downregulated genes unique to B73 were enriched in a network of GO terms related to chlorophyll and metabolic products (right). C, DEGs shared between B73 and *d1* hormone treatments were enriched for a network of GO terms related to gibberellic acid, jasmonic acid and response to abiotic factors in addition to chlorophyll and metabolic products (right). D, Upregulated genes unique to *d1* were enriched for a network of GO terms related to floral meristem determinacy and development (left), downregulated genes unique to *d1* were enriched for a network of GO terms related to chloroplast function and metabolic products (right).

GO term analysis showed that down-regulated DEGs unique to B73 54DAG leaf GA-response are enriched for GO terms related to chlorophyll and metabolic products (Fig. 2B). Of the DEGs that are shared by both B73 and *d1* leaf tissues in response to GA, we saw enrichment of GO terms related to gibberellic acid, jasmonic acid and response to abiotic factors in addition to chlorophyll and metabolic products (Fig. 2C). Further analysis of DEGs that are oppositely regulated in B73 and *d1* plants in response to GA identified *Zm00001eb384740*, an uncharacterized gene associated with RNA polymerase activity as being up-regulated in B73 and down-regulated in d1 during GA treatment. Two other genes, *Zm00001eb267510* (*ZmUMC2730*) and *Zm00001eb040280* (*ZmCKX10*) are down-regulated in B73 and up- regulated in *d1*, and both associated with cytokinin activity (Table 2).

**Table 2.** Significant DEGs that are discordantly regulated in B73 and d1.

| Up-regulated in WT, down-regulated in *d1* |  |  |  |  |  |  |
| --- | --- | --- | --- | --- | --- | --- |
| ID | Description | Gene Ratio | Bg Ratio | p.adjust | Gene ID | Gene symbol |
| GO:0003899 | DNA-directed 5'-3' RNA polymerase activity | 1/1 | 46/39755 | 0.007459 | Zm00001eb384740 | N/A |
| GO:0034062 | 5'-3' RNA polymerase activity | 1/1 | 156/39755 | 0.007459 | Zm00001eb384740 | N/A |
| GO:0097747 | RNA polymerase activity | 1/1 | 157/39755 | 0.007459 | Zm00001eb384740 | N/A |
| Down-regulated in WT, up-regulated in *d1* |  |  |  |  |  |  |
| GO:0005126 | cytokine receptor binding | 1/4 | 10/34321 | 0.007338 | Zm00001eb267510 | umc2730 |
| GO:0019139 | cytokinin dehydrogenase activity | 1/4 | 14/34321 | 0.007338 | Zm00001eb040280 | ckx10 |

GO terms related to chloroplast and metabolic products were also enriched amongst down- regulated DEGs in *d1* leaves in response to GA treatment (Fig. 2D). Finally, while no GO enrichment was detected for up-regulated DEGs in B73 leaf in response to GA, we observed enrichment of GO terms related to development, organ formation, morphogenesis and floral meristem determinacy in DEGs up-regulated in *d1* leaf in response to GA (Fig. 2A,D).

We queried our dataset for specific GA pathway genes and found 293 genes with associated GO terms: gibberellin mediated signaling pathways, response to gibberellin, and gibberellin metabolism (Fig. 3A, Table S5). Of these 293 GA genes, 147/293 (50%) were not DEGs in either B73 or d1 leaf tissues, and 50/293 (17%) were up-regulated or down-regulated in both genotypes, which we refer to as concordantly regulated (Fig. 3B). Numerical fold-change in each maize tissue tested is presented in Supplementary File 2. Amongst GA pathway genes with described functions, *GID1*, which encodes for the GA receptor, was down-regulated in both genotypes (Supplementary File 2). *D8* and *D9*, which encode DELLA proteins involved in repression of GA response, were up-regulated in both B73 and *d1* (Fig. 3C).

**Figure 3.**
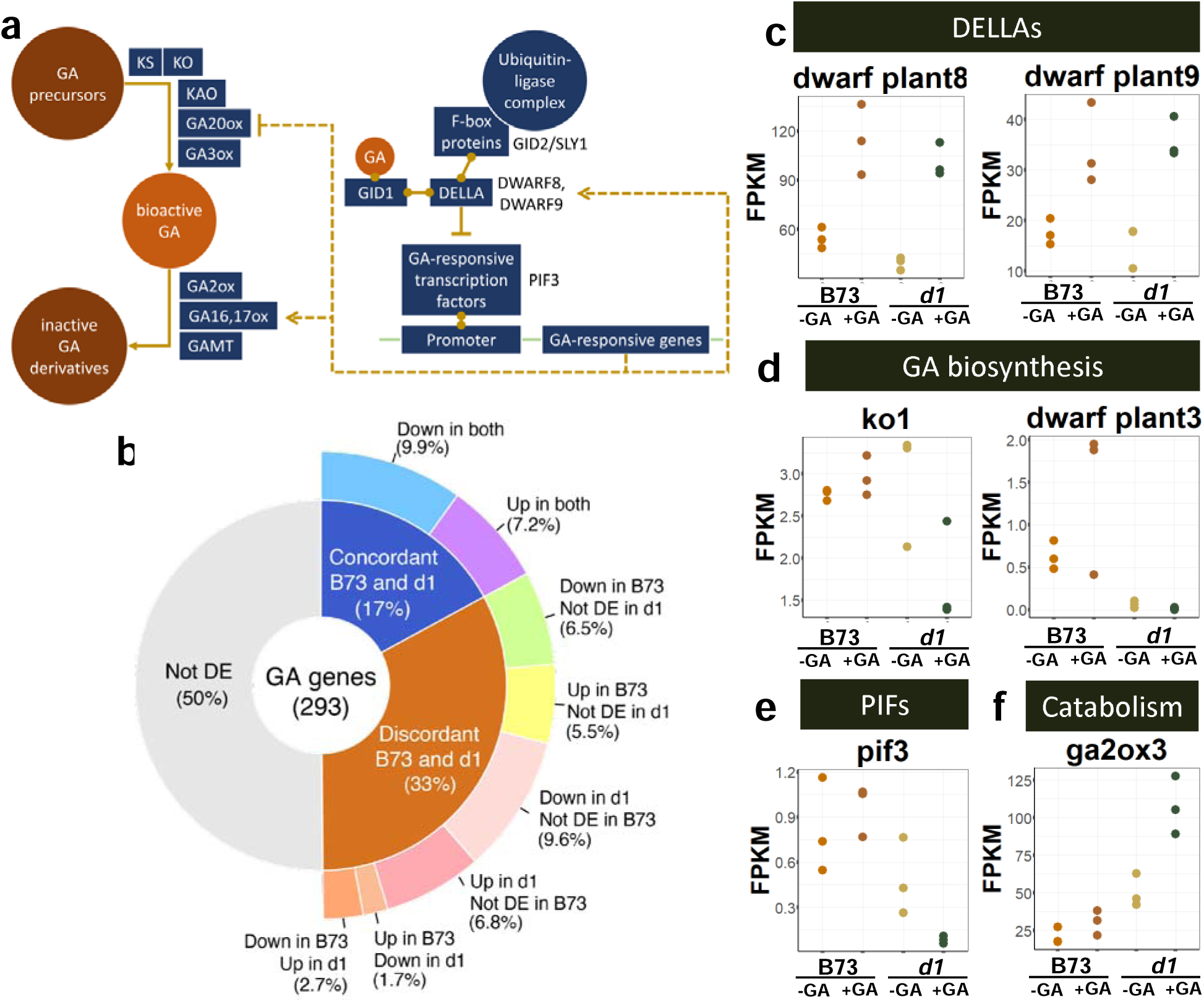
Analysis of GA pathway gene expression reveals widespread discordance between maize B73 and *d1* leaf tissue GA treatment response. A, Known GA biosynthesis and signaling response pathway genes were retrieved using GO terms. B, Of the 293 GA genes detected in our dataset, only 17% displayed concordant differential expression between B73 and *d1* leaves during GA treatment. 33% displayed discordant differential expression between B73 and *d1* leaves during GA treatment. C, Expression of genes from the DELLA family *d8* and *d9* exhibit concordant transcriptional upregulation in B73 and *d1* leaf GA treatment. D, Expression of GA biosynthesis genes *ko1* and *d3* are upregulated in B73 but unchanged in *d1* leaf GA treatments. E, Expression of GA-responsive transcription factor *pif3* is unchanged in B73 but downregulated in *d1* leaf GA treatments. F, Expression of GA-catabolism gene, *ga2ox3* is unchanged in B73 but upregulated in *d1* leaf GA treatments.

The remaining 96/293 GA genes (33%) were differentially regulated in response to GA3 treatment in B73 vs. *d1* leaf tissues, which we refer to as discordantly regulated (Fig. 3B). Amongst these, we found two F-box proteins involved in GA signaling, *GID3* and *GID4* were down-regulated in B73 but not DE in *d1* (Supplementary File 2). Two genes involved in GA biosynthesis, *KAURENE OXIDASE1* (*ZmKO1*) and *DWARF PLANT3* (*ZmD3*) were either up- regulated or not differentially expressed in B73 but down-regulated in *d1* leaf in response to GA (Fig. 3D). Other GA biosynthetic genes such as *GA20ox* family were either not DE or down- regulated in both genotypes (Supplementary File 2). We saw that *PIF3*, a major transcription factor involved in GA response, was not DE in B73 (log2FC = 0.25), but strongly down- regulated in *d1* leaf (log2FC = -2.53) (Fig. 3E). Among genes involved in GA catabolism, *GA2ox3* and *GA2ox6* was not DE in B73 (log2FC = 0.55 and 0.28 respectively) but down- regulated in *d1* (log2FC = 1.09 and 0.61 respectively), while *GA2ox10* was up-regulated in B73 (log2FC = 1.90) and not DE in d1 (log2FC = -0.27) (Fig. 3F, Supplemental File 2). Other *GA2ox* genes were either not DE or up-regulated in both genotypes in response to GA. Other GA genes that were discordantly regulated between B73 and *d1* also include the GRAS transcription factor *SCARECROW LIKE1* (*ZmSCL1*, log2FC B73 = -1.30, log2FC d1 = 1.09) and the *TEOSINTE BRANCHED1, CYCLOIDEA, PCF* (*TCP*) transcription factor *ZmTCPFT26* (log2FC B73 = 0.95, log2FC d1 = -1.25) (Supplementary File 2). Our results suggest that there are major differences in how wild-type and *d1* mutant plants respond to short-term GA treatment, suggesting a fundamental rewiring of GA transcriptional circuitry in the absence of GA biosynthesis.

### RNA-seq analysis between different maize tissue types reveal core GA response machinery

We compared the transcriptomic GA3 treatment responses in organ primordia from different maize tissue types and ages: B73 54DAG leaf, d1 54DAG leaf, B73 90DAG leaf, and B73 90DAG florets and found that B73 90DAG leaf had the largest number of DEGs (2013) and B73 90DAG floret had the smallest (376), while B73 54DAG and d1 54DAG leaves had intermediate DEG numbers (1213 and 1260, respectively; Fig. 4A). Interestingly, B73 90DAG leaf shared relatively few GA-responsive DEGs with B73 54DAG leaf (353 DEGs, corresponding to 29% of DEGs in B73 54DAG leaf, and 18% of those in B73 90DAG leaf), suggesting major changes in how leaf tissues respond to GA treatment as the plant matures. GO-term analysis of DEGs specific to B73 90DAG leaf tissues revealed enrichment of terms related to metabolic processes but not chlorophyll/chloroplast (Fig. 4B). No GO-term enrichment was detected in DEGs specific to B73 90DAG florets (data not shown).

**Figure 4.**
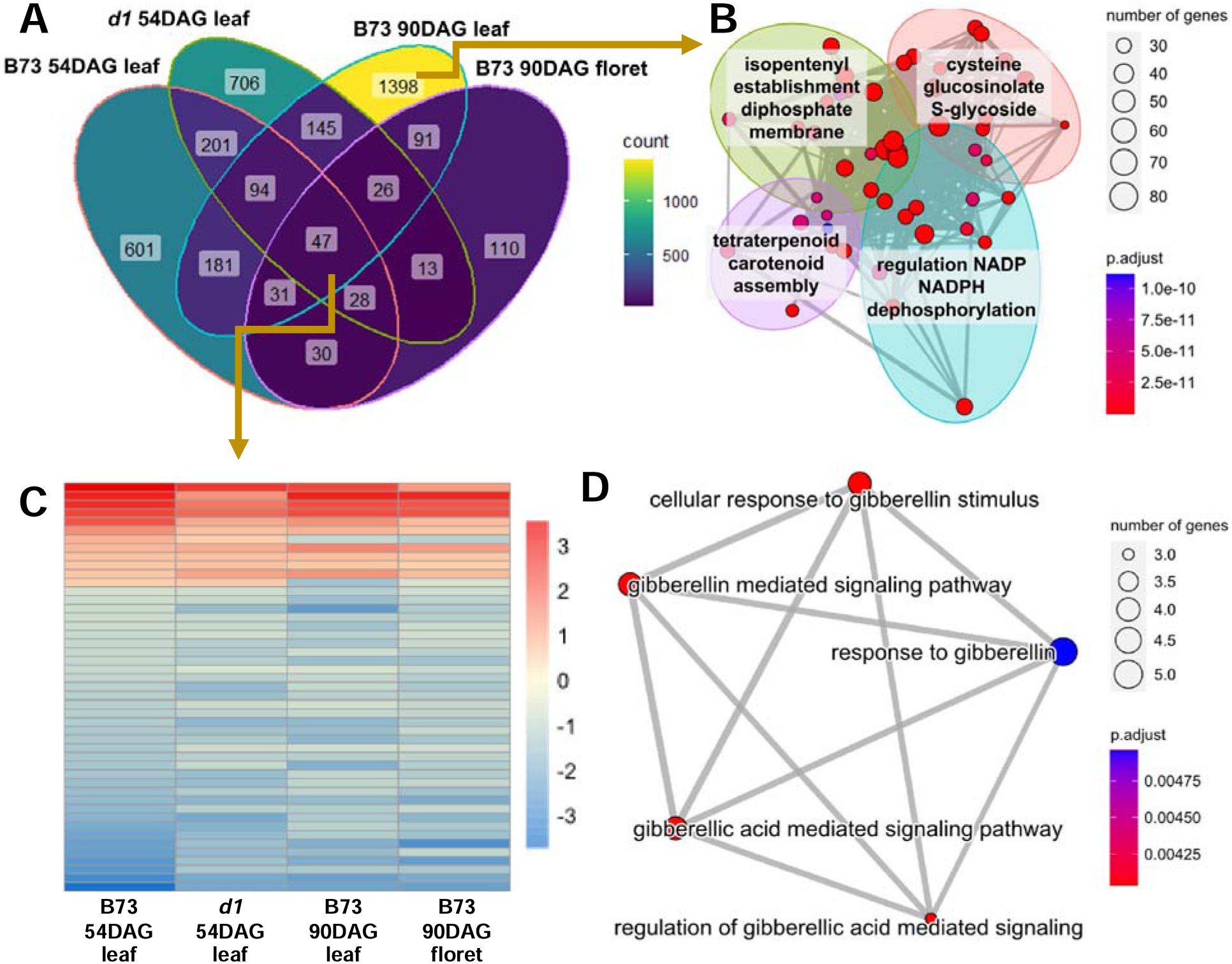
Cross-tissue and cross-genotype G treatment response identifies a core set of differentially regulated maize GA response genes that are independent of GA biosynthesis. A, Venn diagram of B73 54 DAG leaf, d1 54 DAG leaf, B73 90 DAG leaf, and B73 90 DAG floret GA response DEGs with 3,702 total GA-responsive genes across samples. A high proportion of GA-responsive genes were sample-specific. B, DEGs unique to B73 90 DAG leaf were enriched for a network of GO terms related to metabolic products but not photosynthesis, as detected in younger leaf tissue. C, A core set of GA-treatment 47 DEGs were shared by all maize tissues and genotypes. Red, positive log2FC. Blue, negative log2FC. D, The core GA-treatment DEGs were enriched for an interconnected network of GA-related GO terms.

Only 47 genes, representing 1% of the total DEGs, were differentially expressed in response to GA treatment across all four tissues and genotypes (Fig. 4A,C). This group of genes was enriched for GO terms ‘response to gibberellin’ and ‘gibberellin signaling regulation’ and were comprised by core GA response machinery (Fig. 4D, Table 3) and included genes encoding *GID1*, the *GRAS* transcription factors *GRAS2* and *GRAS4*, the *AP2/ERF* transcription factor *EREB2*, and the cell wall-associated protein *ALPHA EXPANSIN1* (*EXPA1*). The full list of core GA-responsive genes is presented in Supplementary File 3, which also includes known F-box proteins such as *GID2* and *FBL41*, another member of the expansin family *GPM282*, proteins involved in sugar signaling and metabolism *HEX5/HXK5* and *AKIN2/SnRK1.1*, as well as proteins in the *AP2/EREBP*, *WRKY*, *MYB* transcription factor, and *zinc finger protein* families.

**Table 3.** GO enrichment of GA-responsive DEGs shared by all tissue types tested.

| geneID | ID | Description | Gene<br>Ratio | Bg<br>Ratio | p.adjust | qvalue |
| --- | --- | --- | --- | --- | --- | --- |
| gras4; gid1;<br>gras2 | GO:0009937 | regulation of gibberellic<br>acid mediated signaling<br>pathway | 3/47 | 46/39755 | 0.004035 | 0.003034 |
| gras4; ereb72;<br>gid1; gras2 | GO:0009740 | gibberellic acid mediated<br>signaling pathway | 4/47 | 156/39755 | 0.004035 | 0.003034 |
| gras4; ereb72;<br>gid1; gras2 | GO:0010476 | gibberellin mediated<br>signaling pathway | 4/47 | 157/39755 | 0.004035 | 0.003034 |
| gras4; ereb72;<br>gid1; gras2 | GO:0071370 | cellular response to<br>gibberellin stimulus | 4/47 | 159/39755 | 0.004035 | 0.003034 |
| gras4; expa1;<br>ereb72; gid1;<br>gras2 | GO:0009739 | response to gibberellin | 5/47 | 351/39755 | 0.004955 | 0.003726 |

### RNA-seq analysis between maize B73 (90DAG) and barley GP (45DAG) leaves and florets

We expanded our transcriptomic analysis of GA response to include a comparison with barley tissues: barley GP 45DAG leaf and barley GP 45DAG floret. We first performed phylogenetic orthology inference using OrthoFinder on the maize, barley, rice, and *Arabidopsis* reference genomes (Emms and Kelly, 2019). 89% and 94% of maize and barley genes were assigned to orthogroups, respectively. 18% of the assigned maize genes and 22.5% of the assigned barley genes were deemed species-specific (i.e, they did not share an orthogroup with genes from any other species). Out of 28,960 orthogroups found, 14,716 (50.8%) contained genes from both maize and barley genomes, while 9,311 (32.2%) contained genes from all four species.

Orthologs were classified based on their phylogenetic relationships, including many-to-many, many-to-one, one-to-many, and one-to-one (Fig. 5A) (Emms and Kelly, 2019). 9,719 maize genes and 6,466 barley genes were found to be in many-to-many relationship, while 4820 maize and barley genes were found to be in one-to-one relationships with each other (Supplemental File 4, Supplemental File 5).

**Figure 5.**
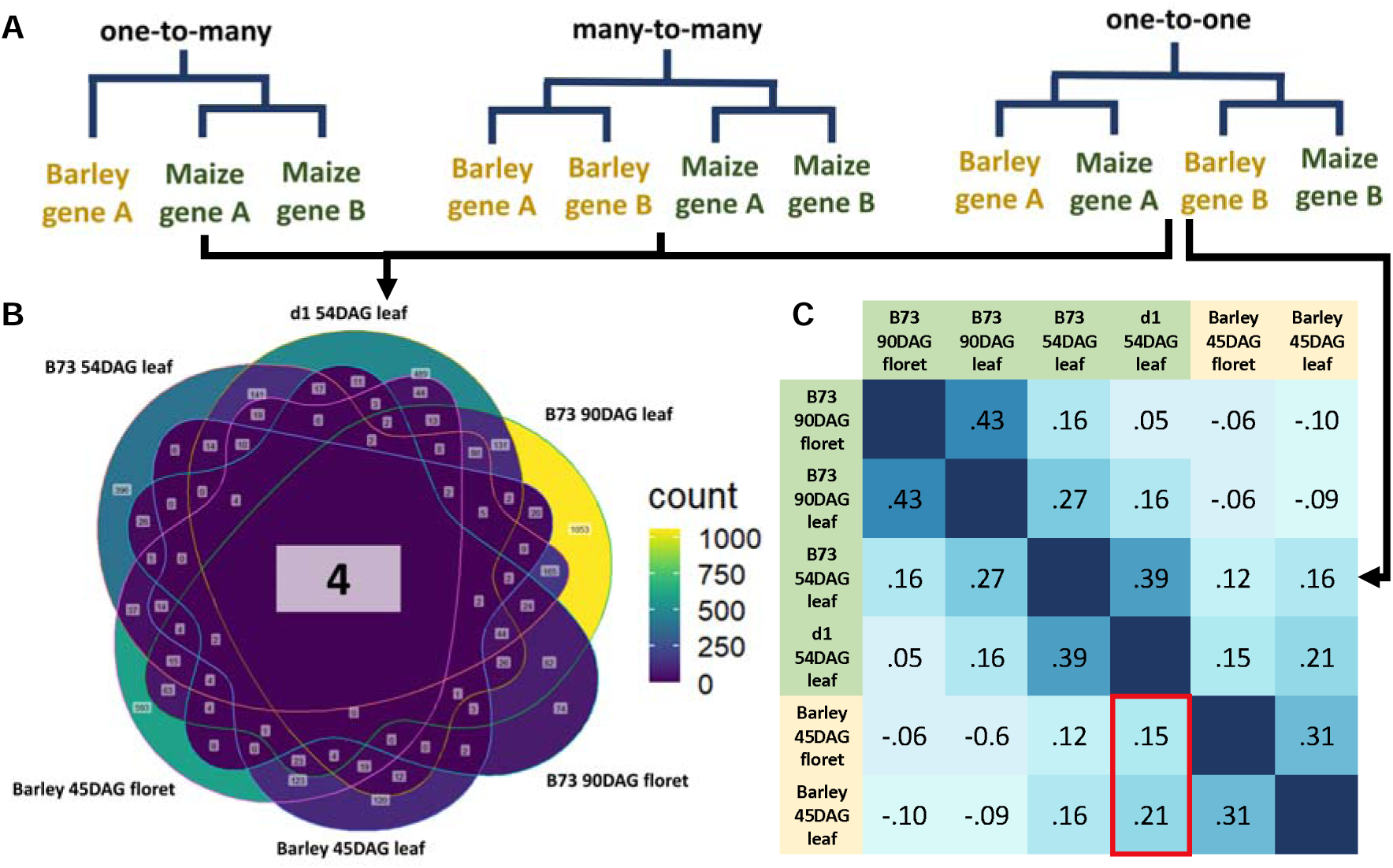
Gene orthology identifies a core orthogroup of GA-response genes while correlation analysis of one-to-one orthologs reveal greater GA-response similarity between maize d1 mutant leaf tissue and barley floral and leaf samples. A, Othogroups were defined as one-to-many, many-to-many, or one-to-one based on Orthofinder phylogenetic homology inference. B, Six-way Venn diagram showing GA treatment differential expression. High numbers of tissue-specific orthogroups were differentially expressed in response to GA treatment (outer rings), while only 4 orthogroups were differentially expressed during GA treatment in all tissues, species, and genotypes examined (center). C, Correlation matrix of one- to-one orthogroup GA-response shows strong similarities based on age, species, and tissue. In cross-species comparison, both barley tissues show greater similarity to vegetative maize d1 tissues than corresponding post flowering tissues from maize B73 (red box). Correlation = Spearman’s ρ.

We matched each DEG with its corresponding orthogroup and looked for overlapping GA-responsive orthogroups among different tissue types in maize and barley. The number of unique orthogroups that contain GA-responsive DEGs for each species and tissue types are presented in Supplementary File 4. Of these GA-responsive orthogroups, only four are shared by all maize and barley tissues tested (Fig. 5B). These orthogroups contain the maize genes encoding for *GID1*, *EXPA1*, *ZINC FINGER PROTEIN7* (*ZNF7*), and *EXPANSIN-B4* (*GPM282*), and their corresponding uncharacterized barley orthologs (Table 4). These genes thus represent the conserved elements of GA signaling and response between maize and barley.

**Table 4.**
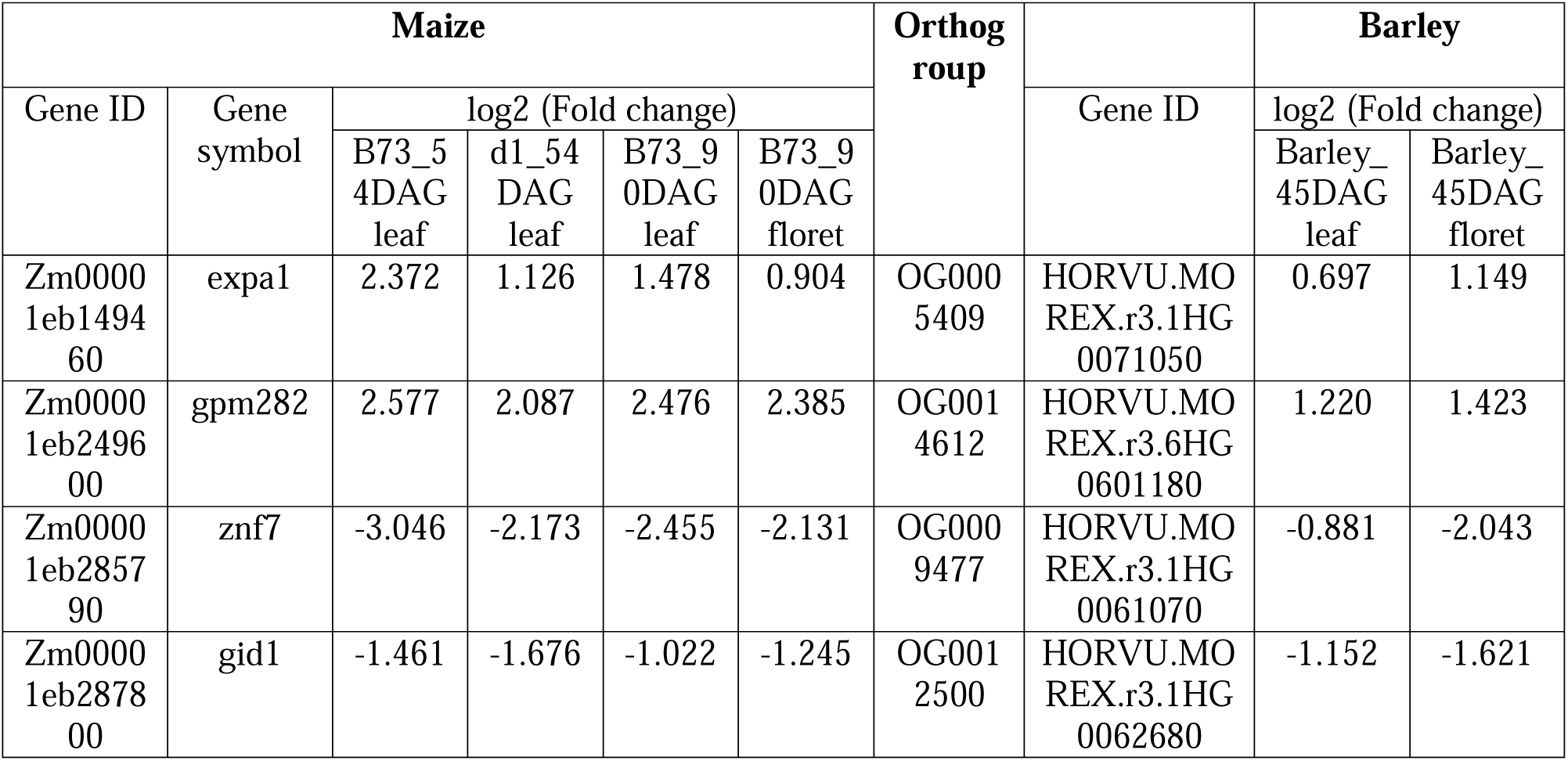
GA-responsive maize and barley gene orthologs.

To eliminate the potential effect of gene subfunctionalization, we focused on one-to-one orthologs between maize and barley to construct a tissue-level correlation matrix of transcriptomic responses to GA3 treatment (Fig. 5C). We found the strongest correlation in one-to-one ortholog GA response between tissues of the same species and age, suggesting that our orthology alignment was robust (maize B73 90DAG leaf vs. B73 90DAG floret - Spearman ρ = 0.43, maize B73 54DAG leaf vs. *d1* 54DAG leaf - ρ = 0.39, barley GP 45DAG leaf vs. GP 45DAG floret - ρ = 0.31) (Fig. 5C). Weak correlations were observed between maize B73 90DAG leaf with either B73 54DAG leaf (ρ = 0.27) or d1 54DAG leaf (ρ = 0.16). We also observed weak correlations between barley 45DAG leaf and floret tissues with maize B73 and d1 54DAG leaf tissues (ρ = 0.12, 0.21 respectively), whereas no correlation was observed between barley 45DAG tissues with maize 90DAG tissues (|ρ| < 0.1, Fig. 5C). Unexpectedly, barley 45DAG leaf and floret both had the greatest correlation with maize *d1* 54DAG leaf tissue in comparison with B73 54DAG leaf tissue, B73 90DAG leaf tissue, or B73 90DAG floral tissue.

### Bombardment and functional validation of DR5ratio and R2D2 auxin sensors in maize and barley

In addition to GA, auxin is another important phytohormone involved in cell identity and organ patterning (Vernoux et al. 2011). We tested the function of DR5v2/DR5 and R2D2 as ratiometric biosensors for auxin in grass tissues using particle bombardment and exogenous hormone application. The DR5v2/DR5 construct contains a DR5v2:ntdTomato and a DR5:n3GFP reporter, both of which are transcriptional reporters that are activated by auxin. DR5v2 was previously demonstrated to show higher sensitivity towards auxin than DR5 (Liao et al. 2015).

The R2D2 construct contains a DII:Venus and mDII:ntdTomato reporter. In the presence of auxin, DII:Venus undergoes protease-mediated degradation, but mDII:ntdTomato is resistant to auxin-dependent degradation (Liao et al. 2015).

The DR5v2/DR5 construct was successfully expressed in maize and barley leaf tissues via particle bombardment (Fig. 6A). We treated bombarded tissues either with 0uM or 10uM IAA over 16 hours, and observed significantly higher DR5v2:ntdTomato fluorescence intensity in IAA-treated compared to mock-treated leaf tissues of both maize and barley (Fig. 6B). In contrast, DR5:n3GFP showed an increase in fluorescence intensity in IAA-treated maize leaf tissues but not barley leaf tissues (Fig. 6B, Table 6). DR5v2 therefore was responsive to exogenous auxin treatment in both maize and barley leaf, and DR5 was only responsive to auxin application in maize leaf but not barley.

**Figure 6.**
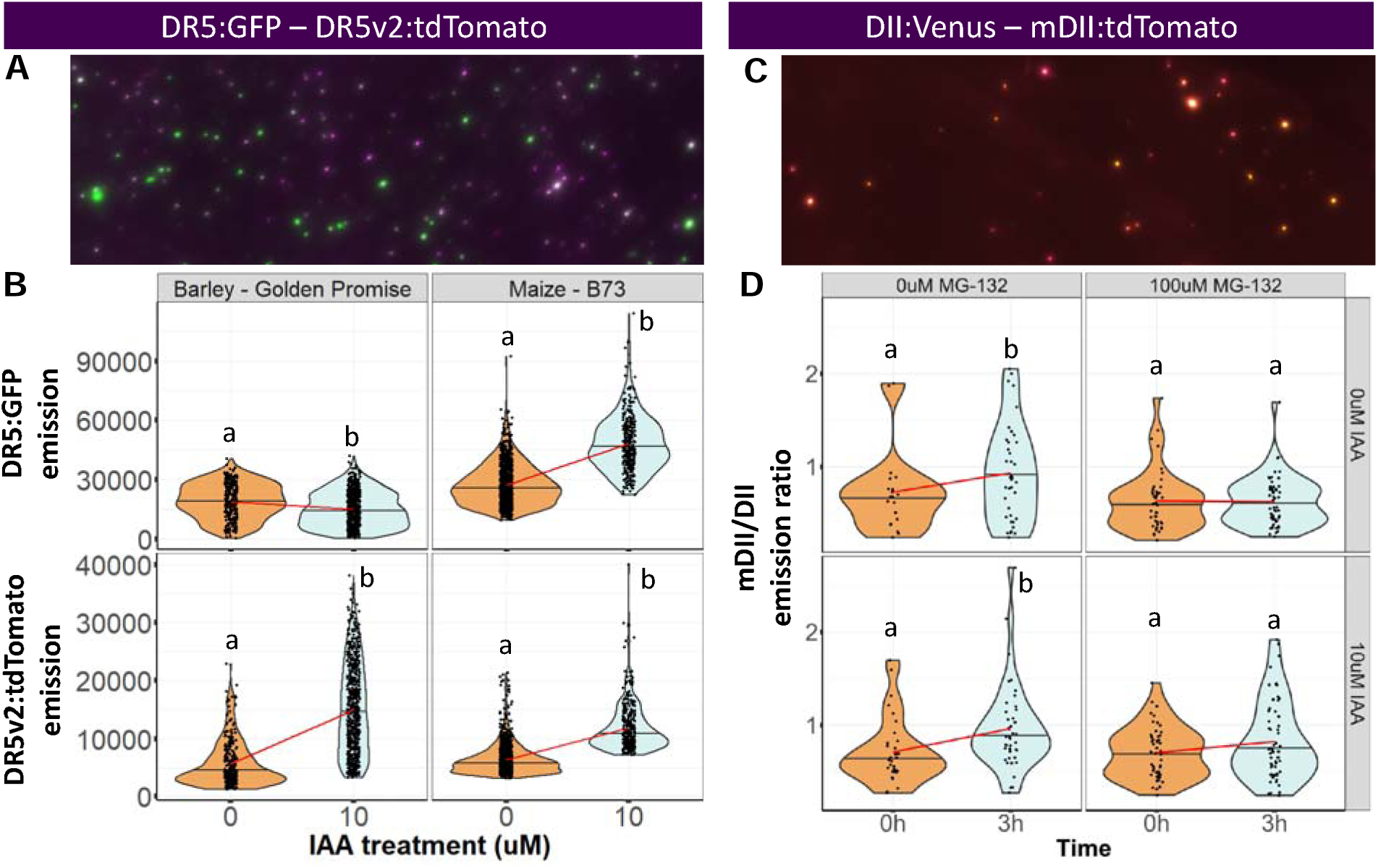
DR5ratio and R2D2 auxin sensors can be expressed in maize and although DR5 is non-responsive, DR5v2 is auxin inducible in barley. A, Representative image of maize leaf tissue bombarded with DR5ratio construct. DR5:GFP, green. DR5v2:tdTomato, magenta. Overlapping signal, white. B, Treatment of bombarded tissues with 0 uM IAA or 10 uM IAA for 24 hours reveals no change in barley DR5-GFP, whereas barley DR5v2:tdTomato and both constructs in maize showed increased fluorescence in 10 uM IAA treatments. Letters, ANOVA posthoc test groups, Emission ∼ IAA treatment, p-value<0.05. C, Representative image of maize tissue bombarded with R2D2 construct. DII:Venus, yellow. mDII:tdTomato, magenta. Overlapping signal, orange. D, Treatment of bombarded tissues with 0 uM IAA or 10 uM IAA for 3 hours reveals increase in mDII/DII emission ratio in maize. Adding 100 uM 26S protease inhibitor, MG-132 leads to no change in mDII/DII emission ratio in maize with 0 uM IAA or 10 uM IAA for 3 hours. Letters, ANOVA posthoc test groups, Emission ratio ∼ IAA treatment, p- value<0.05.

We also observed transient transfection of maize leaf tissues bombarded with the R2D2 construct (Fig. 6C). Tissues were subsequently treated with a combination of 0uM or 10uM IAA with 0uM or 100uM MG-132, which acts as a protease inhibitor. We observed a small but non-significant increase in the mDII/DII emission ratio in maize leaf tissues treated with 0uM IAA and 0uM MG-132 for 3 hours. By contrast, the mDII/DII emission ratio increased significantly in tissues treated with 10uM IAA and 0uM MG-132 after the same time period. Tissues treated either with 0uM IAA and 100uM MG-132 or 10uM IAA and 100uM MG-132 did not show any increase in mDII/DII emission ratio (Fig. 6D, Table 6). These results indicate that R2D2 was functional in maize tissues and was responsive to auxin induction in a protease-dependent manner.

## DISCUSSION

Phytohormones play a ubiquitous role in coordinating plant development and response to biotic and abiotic stimuli. The ability to detect hormone accumulation dynamics at the cellular level has revealed crucial roles of phytohormones such as auxin and cytokinin during stem cell maintenance and tissue patterning (Wang et al. 2014, Wang et al. 2017). Among the phytohormones identified to date, gibberellins are of agricultural interest as they influence plant stature, flowering time, fruit production, and seed germination (Fu et al. 2002, Peng et al. 1999, Sasaki et al., 2002, Ueguchi-Tanaka et al., 2005). A complete mechanistic understanding of GA biology at the cellular and tissue level requires a molecular biosensor that is capable of immediately responding to fluctuations in GA concentration. Such a sensor has yet to be developed in cereal grain models due to high cost and long wait time associated with generating stable transgenic lines.

In this study, we utilized biolistics to generate transient transgenic tissues and validated the GPS2 GA sensor in a variety of models and tissue types. GPS2 revealed differential GA accumulation amongst epidermal cells when comparing maize wild-type B73 and GA-deficient mutant *d1*. Our RNA-seq analysis further suggests a fundamental transcriptional difference in GA response between maize B73 and *d1* plants and revealed core genes involved in GA response in maize and barley. We also tested two new auxin biosensors, DR5v2/DR5 and R2D2 in maize and barley leaf tissues, thus establishing the transient transfection system as a useful approach for functional validation of constructs and comparative study of hormone biology in a short amount of time.

### GPS2 is a functional biosensor for GA, DR5v2 and R2D2 are functional biosensors for auxin in grasses

The RGA-GFP fusion protein used the GA3-inducible degradation of a protein localized to the nucleus as a GA sensor in *Arabidopsis* seedlings (Silverstone et al. 2001). The ability of RGA-GFP to sense GA relies on protease-mediated degradation and no protease-resistant control version has been reported. More recently, GPS1 was developed as a FRET-based optogenetic biosensor, along with its insensitive negative control GPS1-NR both driven by constitutive promoters as sensing is largely independent of sensor levels. GPS1 allows for direct, high-resolution quantification of GA concentration in *Arabidopsis thaliana* (Rizza et al. 2017). We transiently transfected a new version of GPS1, termed GPS2 under the control of the maize UBI promoter, into a variety of grass tissues including leaves and florets from maize, sorghum, barley, and wheat using particle bombardment. GPS2 was successfully expressed in all tissues tested and showed robust increase in FRET emission ratio in response to exogenous applications of GA3. We also observed small but statistically significant increases in the emission ratio of GPS2-NR under exogenous GA treatment, suggesting that GPS2-NR is reporting a response independent of direct GA binding or that optical properties of our tissues are detectably influenced by GA treatment itself. Our observations are consistent with previous reports that showed purified GPS2-NR does not respond to 10 µM GA4 treatment in vitro, but detected small variations in GPS2-NR emission ratio along the *Arabidopsis* root growth axis which are not a direct result of GA accumulation (Rizza et al. 2017).

We additionally tested the DR5v2/DR5 and R2D2 constructs in maize and barley tissues to demonstrate their effectiveness in sensing auxin in grass models, as well as the efficiency of particle bombardment as a functional validation system. DR5v2 and DR5 are transcriptional reporters containing consensus AUXIN RESPONSE FACTOR (ARF) binding sites that are activated in the presence of auxin. The DR5v2/DR5 ratiometric sensor has been tested in *Arabidopsis*, where DR5v2 showed higher level of sensitivity and was able to detect auxin in tissues that DR5 cannot (Liao et al. 2015). We observed expression of the construct in bombarded leaf tissues of maize and barley (Fig. 6A). In maize leaf tissue, both DR5v2:ntdTomato and DR5:n3GFP showed increases in signal intensity in response to IAA treatment. In contrast, only DR5v2 increased in signal intensity in response to IAA in barley leaf tissue (Fig. 6B, Table 5). Our observations thus support DR5v2 as the more effective auxin sensor in a broad range of grass tissues.

**Table 5.** DR5/DR5v2 emission intensity in response to IAA treatment in barley and maize.

| Species | IAA (uM) | N | DR5:n3GFP | ci | DR5v2:ntdTomato | ci |
| --- | --- | --- | --- | --- | --- | --- |
| **Barley – Golden Promise** | 0uM | 294 | 19099.17 | 998.92 | 5634.87 | 228.4 |
|  | 10uM | 658 | 15051.71 | 669.65 | 14966.31 | 302.48 |
| **Maize – B73** | 0uM | 1088 | 27348.75 | 605.34 | 6413.37 | 156.23 |
|  | 10uM | 328 | 48123.40 | 1496.81 | 11848.93 | 443.79 |

The R2D2 ratiometric reporter carries the DII-Venus fusion protein, which undergoes auxin-dependent degradation, as well as the auxin-resistant mDII-ntdTomato, both under the control of the Arabidopsis RIBOSOMAL PROTEIN 5A (AtRPS5AS) promoter (Liao et al. 2015). The construct was expressed by maize leaf tissue in transient transfection and showed increased mDII/DII emission ratio in response to IAA treatment (Fig. 6C,D, Table 6). The increase in emission ratio was abolished in treatment with both IAA and the protease inhibitor MG-132 as previously reported in *Arabidopsis* (Liao et al. 2015), suggesting that R2D2’s response was specific to IAA and IAA-mediated degradation. Therefore, R2D2 is an effective sensor for auxin in maize leaf, and its function in other species/tissue types remains to be investigated.

**Table 6.** R2D2 emission ratio in response to IAA/MG-132 treatments in maize.

| IAA (uM) | MG-132 (uM) | Time | N | mDII.DII | ci | Cliff's delta |
| --- | --- | --- | --- | --- | --- | --- |
| 0uM | 0uM | 0h | 20 | 0.726894 | 0.208453 | - |
| 0uM | 0uM | 3h | 38 | 0.933582 | 0.169989 | 0.28 |
| 0uM | 100uM | 0h | 43 | 0.632518 | 0.098871 | - |
| 0uM | 100uM | 3h | 56 | 0.622163 | 0.069831 | 0.06 |
| 10uM | 0uM | 0h | 33 | 0.711184 | 0.124407 | - |
| 10uM | 0uM | 3h | 42 | 0.964243 | 0.143913 | 0.41 |
| 10uM | 100uM | 0h | 50 | 0.704288 | 0.077689 | - |
| 10uM | 100uM | 3h | 61 | 0.824499 | 0.106809 | 0.12 |

### B73 and *d1* transcriptomes demonstrate rewiring of downstream GA-response

Treatment of tissues transiently expressing GPS2 with exogenous GA3 revealed a difference in the regulation of GA accumulation between B73 and *d1* mutant. The *D1* gene encodes a GA3ox that catalyzes the final steps of bioactive GA synthesis, and the *d1* loss-of-function mutant accumulates lower levels of GA1/GA5 and higher levels of GA20/GA29 than the wild type (Chen et al. 2014, Fujioka et al. 1988, Spray et al. 1996). The dwarfism phenotype of *d1* seedlings can be rescued with application of 10uM GA3 for 7 days (Chen et al. 2014). Due to the low endogenous level of bioactive GA in *d1*, we had expected GA3 treatment to produce a more pronounced increase in GPS2 emission ratio in *d1* than in B73 leaf tissues. Surprisingly, we observed a reduction in GPS2 emission ratio in *d1* leaf treated with GA3 for 16h amongst the epidermal nuclei we observed by bombardment. Given the low affinity of GPS to non-bioactive GAs, our results suggest that the endogenous level of bioactive GA of *d1* is decreased when exogenous GA is applied (Rizza et al. 2017). A temporary increase in GA concentration is known to trigger negative feedback mechanisms to maintain cellular GA homeostasis, via the down-regulation of GA biosynthetic genes, and up-regulation of GA catabolic genes (Ogawa et al. 2003). Such a negative feedback mechanism might be responsible for the observed decrease in GPS emission ratio in *d1* leaves to cope with drastic changes in GA levels. Our transient system does not allow the observation of GPS2 emission in all cells within a tissue, and cells measured near the epidermis may experience increased export or decreased GA import despite an overall increase in GA after treatment. There may be nuanced spatial regulation of GA accumulation across cell layers or cell types which would be revealed through the production and analysis of stable GPS2 transformants.

We performed comparative transcriptomic analysis between wild-type and *d1* leaf tissues with either mock or GA3 treatment to explore their potential differences in GA response mechanisms. Overall, we observed large differences in how the B73 and *d1* transcriptomes respond to GA, as only 17% of DEGs were shared by both (Fig. 2A). GO analysis of DEGs reveal major changes in metabolic pathways in response to GA induction in both genotypes. In addition, development and morphogenesis-related GO terms were specifically enriched in DEGs of *d1* mutants (Fig. 2D). These data suggest that *d1* plants are fundamentally different from wild-type at the transcriptional level, possibly to cope with the lifelong, global deficiency in bioactive GAs, and thus might undergo major developmental reprogramming after GA treatment.

Among the genes involved in GA metabolism and signaling that are differentially regulated by GA3 treatment, the majority are regulated in different directions between B73 and *d1* (Fig. 3B). We found down-regulation of the gene encoding GA receptor GID1, as well as up-regulation of genes encoding the GA repressors DWARF8 and DWARF9 as indication of GA negative feedback were shared across B73 and *d1* (Fig. 3C, Supplementary File 2). We observed strong down-regulation of *PIF3* transcript level, a major positive regulator of GA response, in GA-treated *d1* leaves, whereas *PIF3* level was not affected in GA-treated wild-type leaves (Fig. 3E). Other members of the PIF family have been shown to participate in an auto-regulatory negative loop during thermomorphogenesis in *Arabidopsis* (Lee et al. 2021). Thus, our observation of *PIF3* downregulation may represent extreme transcriptional feedback after treatment in *d1* leaves compared to wild-type, which maintains constant *PIF3* levels by autoregulation. We also saw down-regulation of GA biosynthetic genes and up-regulation of GA catabolic genes in both B73 and *d1* (Supplementary File 2). Additional GA-metabolic genes were specifically responsive to GA in *d1*, including *KO1*, *D3*, *GA2ox3* and *GA2ox6* (Fig. 3F, Supplementary File 2). Taken together, our RNA-seq data support the presence of a negative feedback mechanism for GA response in both the wild-type and *d1* mutant. However, the negative feedback could be stronger in *d1* plants, possibly as an adaptation to homeostasis at a low level of bioactive GA. Given that long-term treatment with GA3 can restore *d1* to wild-type phenotype, we expect the response of GA-related genes and overall transcriptome of B73 and *d1* to become more similar with repeated treatment beyond the 24-hour endpoint that we report.

### Core elements of GA response within maize, and between maize and barley

By comparing the transcriptomic profiles of different maize and barley tissues, we identified a list of core sequence homology orthogroups involved in GA response that are shared across tissue types and species. In maize-maize comparisons, 47 core GA-responsive genes were shared by leaf and floret tissues of B73 plants at 90DAG, as well as B73 and *d1* plants at 54DAG. Notable genes include *GID1* and *GID2*, both of which are known elements in GA signaling pathway (Davière and Achard, 2013). The core list includes another F-box protein, *ZmFBL41*, which is involved in disease resistance in maize and rice and has not previously been linked to GA response (Li et al. 2019). Our core GA-responsive gene list also demonstrates a conserved link between sugar metabolism and GA signaling, including hexokinase protein family members, ZmHEX5/ZmHKX5, and AMPK/SNF1 protein kinase, ZmAKIN2/ZmSnRK1.1, both of which are conserved in all organisms and have demonstrated roles in sugar metabolism and developmental signaling (Beana-González et al. 2007, Jang et al. 1997, Rolland et al. 2006).

Sugar signaling interacts with GA pathways in embryo development, and ZmHKX5 expression in shoot tissues was shown to be induced by GA treatment (Perata et al. 1997, Zhang et al. 2014). ZmSnRK1 is involved in the trehalose signaling pathway that regulates energy allocation as well as plant growth (Paul et al. 2020), but the link between GA and trehalose signaling pathways has not yet been elucidated. Our core maize GA-inducible list includes *ZmGPM282* and *ZmEXPA1*, which both belong to the expansin family of proteins involved in cell wall loosening during cell growth, elongation, and fertilization (Cosgrove 2000, Penning et al. 2009). GA is a known regulator of cell expansion, and low-phosphate stress affects expression of both expansin genes and GA signaling/biosynthetic genes, demonstrating their potential co-regulation (de Lucas et al. 2008, Zhang et al. 2019). The list of core maize GA-responsive genes also includes members of the GRAS, AP2/EREBP, WRKY, MYB, and zinc finger protein transcription factor families, all of which contain members known to interact with GA signaling (Bolle et al. 2000, Day et al. 2004, Devaiah et al. 2009, Gubler et al. 1995, Ward et al. 2006, Xie et al. 2019, Yaish et al. 2010, Zhang et al. 2004, Zhang et al. 2011, Zhang et al, 2015, Zhou et al. 2013).

In addition, we identified four pairs of orthologs that are differentially regulated in response to GA3 treatment in both maize and barley. The maize orthologs GID1, GPM282, EXPA1, and ZNF7 are also present in our list of 47 core maize-maize GA-responsive genes. Other than GID1, the function of these genes in relation to GA signaling have not been characterized in maize or barley. Thus, they present new avenues to explore the conserved role of GA in grasses, as well as candidates for precision breeding to improve crop yield and architecture.

### Barley Golden Promise, a G Protein γ Subunit mutant, shares similarities with maize *d1* GA biosynthesis mutant

Our analysis of 1-to-1 orthologs shows that the transcriptomic GA-response of barley Golden Promise leaves and floral tissues is more similar to the leaf GA-response from maize *d1,* a *GA3ox2* GA biosynthesis mutant, than to the GA-response of wild-type maize B73 leaf or floral tissues. Modern elite varieties of barley, rice, wheat, and several other row crops were selected for a semi-dwarf growth habit during the Green Revolution. Semi-dwarf varieties of barley with *sdw1*/*denso* alleles possess mutations in *GA20ox2*, a key enzyme in GA biosynthesis (Xu et al. 2017). However, Golden Promise along with several other barley varieties derive semi-dwarf stature from the *ari-e* (*breviaristatum-e*) locus (Kuczyńska et al. 2013). The Golden Promise *ari- e.GP* allele is GA-insensitive, suggesting that its mechanism of dwarfing involves decreased GA signaling (Pakniyat et al. 2006, Foster 2002). Yet, the *ari-e.GP* locus results from a loss-of-function mutation in *HvDep1*, a G Protein γ Subunit, part of the heterotrimeric G protein complex (Wendt et al. 2016). Although in rice G Protein α Subunit function is necessary for GA perception (Ueguchi-Tanaka et al. 2000), a link between G Protein γ Subunit and GA signaling has not been established. Work with barley protoplasts has shown that GA can be perceived at the plasma membrane (Gilroy and Jones, 1994), which would be consistent with a G Protein-mediated GA signaling mechanism that is spatially separated from nuclear GID1-mediated GA perception. We anticipate that barley G Protein-mediated GA signaling mediated by the *ari-e* locus must ultimately converge on GID1 targets through a yet unknown mechanism, leading to a similar transcriptional phenotype between maize *d1* and barley Golden Promise during GA treatment.

## AUTHOR CONTRIBUTIONS

Designed experiments: TQD, CD, AMJ, SL. Performed experiments: TQD. Analyzed data: TQD. Prepared figures: TQD. Designed project: TQD, SL. Wrote and revised manuscript: TQD, CD, AMJ, SL. Secured funding: SL, CD, AMJ.

## ACKNOWLEDGEMENTS

We thank Sarah Hake for kindly providing us with the *dwarf1* mutant. We thank Zuzana Vejlupkova for particle bombardment assistance, and Jeff Bishop for Illumina sequencing assistance. We acknowledge the USDA National Germplasm Resources Laboratory for providing several seed stocks. SL and TQD are supported by NSF IOS-1922543 and IOS- 2211434. AMJ is supported by the Gatsby Charitable Trust (GAT3395). CD is supported by H2020-IF (844398).

## REFERENCES

Baena-González, E.; Rolland, F.; Thevelein, J. M.; Sheen, J. A Central Integrator of Transcription Networks in Plant Stress and Energy Signalling. Nature 2007, 448 (7156), 938–942. 10.1038/nature06069.

Binenbaum, Jenia, Roy Weinstain, and Eilon Shani. 2018. “Gibberellin Localization and Transport in Plants.” Trends in Plant Science 23 (5): 410–21. 10.1016/j.tplants.2018.02.005.

Bolle, C.; Koncz, C.; Chua, N.-H. PAT1, a New Member of the GRAS Family, Is Involved in Phytochrome A Signal Transduction. Genes Dev. 2000, 14 (10), 1269–1278. 10.1101/gad.14.10.1269.

Brian, P. W. Effects of Gibberellins on Plant Growth and Development. Biological Reviews 1959, 34 (1), 37–77. 10.1111/j.1469-185X.1959.tb01301.x.

Carlson M, Pagès H (2022). AnnotationForge: Tools for building SQLite-based annotation data packages. R package version 1.38.0, https://bioconductor.org/packages/AnnotationForge.

Chen, Y.; Hou, M.; Liu, L.; Wu, S.; Shen, Y.; Ishiyama, K.; Kobayashi, M.; McCarty, D. R.; Tan, B.-C. The Maize DWARF1 Encodes a Gibberellin 3-Oxidase and Is Dual Localized to the Nucleus and Cytosol1[W]. Plant Physiol 2014, 166 (4), 2028–2039. 10.1104/pp.114.247486.

Ci, J.; Wang, X.; Wang, Q.; Zhao, F.; Yang, W.; Cui, X.; Jiang, L.; Ren, X.; Yang, W. Genome-Wide Analysis of Gibberellin-Dioxygenases Gene Family and Their Responses to GA Applications in Maize. PLoS ONE 2021, 16 (5), e0250349. 10.1371/journal.pone.0250349.

Cosgrove, D. J. Loosening of Plant Cell Walls by Expansins. Nature 2000, 407 (6802), 321–326. 10.1038/35030000.

Davière, J.-M.; Achard, P. Gibberellin Signaling in Plants. Development 2013, 140 (6), 1147–1151. 10.1242/dev.087650.

Day, R. B.; Tanabe, S.; Koshioka, M.; Mitsui, T.; Itoh, H.; Ueguchi-Tanaka, M.; Matsuoka, M.; Kaku, H.; Shibuya, N.; Minami, E. Two Rice GRAS Family Genes Responsive to N-Acetylchitooligosaccharide Elicitor Are Induced by Phytoactive Gibberellins: Evidence for Cross-Talk Between Elicitor and Gibberellin Signaling in Rice Cells. Plant Mol Biol 2004, 54 (2), 261–272. 10.1023/B:PLAN.0000028792.72343.ee.

de Lucas, M.; Davière, J.-M.; Rodríguez-Falcón, M.; Pontin, M.; Iglesias-Pedraz, J. M.; Lorrain, S.; Fankhauser, C.; Blázquez, M. A.; Titarenko, E.; Prat, S. A Molecular Framework for Light and Gibberellin Control of Cell Elongation. Nature 2008, 451 (7177), 480–484. 10.1038/nature06520.

Devaiah, B. N.; Madhuvanthi, R.; Karthikeyan, A. S.; Raghothama, K. G. Phosphate Starvation Responses and Gibberellic Acid Biosynthesis Are Regulated by the MYB62 Transcription Factor in Arabidopsis. Molecular Plant 2009, 2 (1), 43–58. 10.1093/mp/ssn081.

El-Sharkawy, I.; Sherif, S.; Abdulla, M.; Jayasankar, S. Plum Fruit Development Occurs via Gibberellin–Sensitive and –Insensitive DELLA Repressors. PLOS ONE 2017, 12 (1), e0169440. 10.1371/journal.pone.0169440.

Emms, D. M.; Kelly, S. OrthoFinder: Phylogenetic Orthology Inference for Comparative Genomics. Genome Biology 2019, 20 (1), 238. 10.1186/s13059-019-1832-y.

Engler, C.; Youles, M.; Gruetzner, R.; Ehnert, T.M.; Werner, S.; Jones, J.D.G.; Patron, N.J.; Marillonnet, S. A Golden Gate Modular Cloning Toolbox for Plants. ACS Synthetic Biology 2014, 3 (11): 839–43. 10.1021/sb4001504.

Fleet, C. M.; Sun, T. A DELLAcate Balance: The Role of Gibberellin in Plant Morphogenesis. Current Opinion in Plant Biology 2005, 8 (1), 77–85. 10.1016/j.pbi.2004.11.015.

Forster, Brian P. 2001. “Mutation Genetics of Salt Tolerance in Barley: An Assessment of Golden Promise and Other Semi-Dwarf Mutants.” Euphytica/ Netherlands Journal of Plant Breeding 120 (3): 317–28. 10.1023/A:1017592618298.

Fu, X.; Richards, D. E.; Ait-ali, T.; Hynes, L. W.; Ougham, H.; Peng, J.; Harberd, N. P. Gibberellin-Mediated Proteasome-Dependent Degradation of the Barley DELLA Protein SLN1 Repressor. The Plant Cell 2002, 14 (12), 3191–3200. 10.1105/tpc.006197.

Fujioka, S.; Yamane, H.; Spray, C. R.; Gaskin, P.; Macmillan, J.; Phinney, B. O.; Takahashi, N. Qualitative and Quantitative Analyses of Gibberellins in Vegetative Shoots of Normal, Dwarf-1, Dwarf-2, Dwarf-3, and Dwarf-5 Seedlings of Zea Mays L. 1. Plant Physiol 1988, 88 (4), 1367–1372.

Gollin, D.; Hansen, C. W.; Wingender, A. M. Two Blades of Grass: The Impact of the Green Revolution. Journal of Political Economy 2021. 10.1086/714444.

Gubler, F.; Kalla, R.; Roberts, J. K.; Jacobsen, J. V. Gibberellin-Regulated Expression of a Myb Gene in Barley Aleurone Cells: Evidence for Myb Transactivation of a High-PI Alpha-Amylase Gene Promoter. Plant Cell 1995, 7 (11), 1879–1891.

He, J.; Xin, P.; Ma, X.; Chu, J.; Wang, G. Gibberellin Metabolism in Flowering Plants: An Update and Perspectives. Frontiers in Plant Science 2020, 11.

Helliwell, C. A.; Chandler, P. M.; Poole, A.; Dennis, E. S.; Peacock, W. J. The CYP88A Cytochrome P450, Ent-Kaurenoic Acid Oxidase, Catalyzes Three Steps of the Gibberellin Biosynthesis Pathway. Proceedings of the National Academy of Sciences 2001, 98 (4), 2065–2070. 10.1073/pnas.98.4.2065.

Helliwell, C. A.; Poole, A.; James Peacock, W.; Dennis, E. S. Arabidopsis Ent-Kaurene Oxidase Catalyzes Three Steps of Gibberellin Biosynthesis. Plant Physiology 1999, 119 (2), 507–510. 10.1104/pp.119.2.507.

Hirose, F.; Inagaki, N.; Hanada, A.; Yamaguchi, S.; Kamiya, Y.; Miyao, A.; Hirochika, H.; Takano, M. Cryptochrome and Phytochrome Cooperatively but Independently Reduce Active Gibberellin Content in Rice Seedlings under Light Irradiation. Plant and Cell Physiology 2012, 53 (9), 1570–1582. 10.1093/pcp/pcs097.

Jang, J. C.; León, P.; Zhou, L.; Sheen, J. Hexokinase as a Sugar Sensor in Higher Plants. The Plant Cell 1997, 9 (1), 5–19. 10.1105/tpc.9.1.5.

Jiang, Z.; Xu, G.; Jing, Y.; Tang, W.; Lin, R. Phytochrome B and REVEILLE1/2- Mediated Signalling Controls Seed Dormancy and Germination in Arabidopsis. Nat Commun 2016, 7 (1), 12377. 10.1038/ncomms12377.

Kanno, Yuri, Takaya Oikawa, Yasutaka Chiba, Yasuhiro Ishimaru, Takafumi Shimizu, Naoto Sano, Tomokazu Koshiba, Yuji Kamiya, Minoru Ueda, and Mitsunori Seo. 2016. “AtSWEET13 and AtSWEET14 Regulate Gibberellin-Mediated Physiological Processes.” Nature Communications 7 (October): 13245. 10.1038/ncomms13245.

Kirienko, D. R.; Luo, A.; Sylvester, A. W. Reliable Transient Transformation of Intact Maize Leaf Cells for Functional Genomics and Experimental Study. Plant Physiology 2012, 159 (4), 1309–1318. 10.1104/pp.112.199737.

Kuczyńska, Anetta, Maria Surma, Tadeusz Adamski, Krzysztof Mikołajczak, Karolina Krystkowiak, and Piotr Ogrodowicz. Effects of the Semi-Dwarfing sdw1/denso Gene in Barley. Journal of Applied Genetics 2013 54 (4): 381–90. 10.1007/s13353-013-0165-x.

Lawit, S. J.; Wych, H. M.; Xu, D.; Kundu, S.; Tomes, D. T. Maize DELLA Proteins Dwarf Plant8 and Dwarf Plant9 as Modulators of Plant Development. Plant and Cell Physiology 2010, 51 (11), 1854–1868. 10.1093/pcp/pcq153.

Lee, Sanghwa, Ling Zhu, and Enamul Huq. 2021. “An Autoregulatory Negative Feedback Loop Controls Thermomorphogenesis in Arabidopsis.” PLoS Genetics 17 (6): e1009595. 10.1371/journal.pgen.1009595.

Li, N.; Lin, B.; Wang, H.; Li, X.; Yang, F.; Ding, X.; Yan, J.; Chu, Z. Natural Variation in ZmFBL41 Confers Banded Leaf and Sheath Blight Resistance in Maize. Nat Genet 2019, 51 (10), 1540–1548. 10.1038/s41588-019-0503-y.

Liao, C.-Y.; Smet, W.; Brunoud, G.; Yoshida, S.; Vernoux, T.; Weijers, D. Reporters for Sensitive and Quantitative Measurement of Auxin Response. Nat Methods 2015, 12 (3), 207–210. 10.1038/nmeth.3279.

Mertz, Dan. 1985. “Effect of Phytochrome on Uptake and Efflux of Gibberellin.” Plant & Cell Physiology 26 (4): 701–7. 10.1093/oxfordjournals.pcp.a076960.

Ogawa, M.; Hanada, A.; Yamauchi, Y.; Kuwahara, A.; Kamiya, Y.; Yamaguchi, S. Gibberellin Biosynthesis and Response during Arabidopsis Seed Germination. Plant Cell 2003, 15 (7), 1591–1604. 10.1105/tpc.011650.

Olszewski, N.; Sun, T.; Gubler, F. Gibberellin Signaling. Plant Cell 2002, 14 (Suppl), s61–s80. 10.1105/tpc.010476.

Pakniyat, H., W. T. B. Thomas, P. D. S. Caligari, and B. P. Forster. Comparison of Salt Tolerance of GPert and Non-GPert Barleys. Plant Breeding = Zeitschrift Fur Pflanzenzuchtung 1997 116 (2): 189–91. 10.1111/j.1439-0523.1997.tb02177.x.

Paul, M. J.; Watson, A.; Griffiths, C. A. Trehalose 6-Phosphate Signalling and Impact on Crop Yield. Biochemical Society Transactions 2020, 48 (5), 2127–2137. 10.1042/BST20200286.

Peng, J.; Carol, P.; Richards, D. E.; King, K. E.; Cowling, R. J.; Murphy, G. P.; Harberd, N. P. The Arabidopsis GAI Gene Defines a Signaling Pathway That Negatively Regulates Gibberellin Responses. Genes Dev. 1997, 11 (23), 3194–3205. 10.1101/gad.11.23.3194.

Peng, J.; Richards, D. E.; Hartley, N. M.; Murphy, G. P.; Devos, K. M.; Flintham, J. E.; Beales, J.; Fish, L. J.; Worland, A. J.; Pelica, F.; Sudhakar, D.; Christou, P.; Snape, J. W.; Gale, M. D.; Harberd, N. P. ‘Green Revolution’ Genes Encode Mutant Gibberellin Response Modulators. Nature 1999, 400 (6741), 256–261. 10.1038/22307.

Penning, B. W.; Hunter, C. T., III; Tayengwa, R.; Eveland, A. L.; Dugard, C. K.; Olek, A. T.; Vermerris, W.; Koch, K. E.; McCarty, D. R.; Davis, M. F.; Thomas, S. R.; McCann, M. C.; Carpita, N. C. Genetic Resources for Maize Cell Wall Biology. Plant Physiology 2009, 151 (4), 1703–1728. 10.1104/pp.109.136804.

Perata, P.; Matsukura, C.; Vernieri, P.; Yamaguchi, J. Sugar Repression of a Gibberellin-Dependent Signaling Pathway in Barley Embryos. The Plant Cell 1997, 9 (12), 2197–2208. 10.1105/tpc.9.12.2197.

Rieu, I.; Eriksson, S.; Powers, S. J.; Gong, F.; Griffiths, J.; Woolley, L.; Benlloch, R.; Nilsson, O.; Thomas, S. G.; Hedden, P.; Phillips, A. L. Genetic Analysis Reveals That C19-GA 2-Oxidation Is a Major Gibberellin Inactivation Pathway in Arabidopsis. The Plant Cell 2008, 20 (9), 2420–2436. 10.1105/tpc.108.058818.

Rizza, A.; Walia, A.; Lanquar, V.; Frommer, W. B.; Jones, A. M. In Vivo Gibberellin Gradients Visualized in Rapidly Elongating Tissues. Nature Plants 2017, 3 (10), 803–813.

Rizza, A.; Walia, A.; Tang, B.; Jones, A. M. Visualizing Cellular Gibberellin Levels Using the NlsGPS1 Förster Resonance Energy Transfer (FRET) Biosensor. JoVE 2019, No. 143, 58739. 10.3791/58739.

Rolland, F.; Baena-Gonzalez, E.; Sheen, J. SUGAR SENSING AND SIGNALING IN PLANTS: Conserved and Novel Mechanisms. Annual Review of Plant Biology 2006, 57 (1), 675–709. 10.1146/annurev.arplant.57.032905.105441.

Sasaki, A.; Ashikari, M.; Ueguchi-Tanaka, M.; Itoh, H.; Nishimura, A.; Swapan, D.; Ishiyama, K.; Saito, T.; Kobayashi, M.; Khush, G. S.; Kitano, H.; Matsuoka, M. A Mutant Gibberellin-Synthesis Gene in Rice. Nature 2002, 416 (6882), 701–702. 10.1038/416701a.

Serrani, J. C.; Sanjuán, R.; Ruiz-Rivero, O.; Fos, M.; García-Martínez, J. L. Gibberellin Regulation of Fruit Set and Growth in Tomato. Plant Physiol 2007, 145 (1), 246–257. 10.1104/pp.107.098335.

Shani, E.; Weinstain, R.; Zhang, Y.; Castillejo, C.; Kaiserli, E.; Chory, J.; Tsien, R. Y.; Estelle, M. Gibberellins Accumulate in the Elongating Endodermal Cells of Arabidopsis Root. Proceedings of the National Academy of Sciences 2013, 110 (12), 4834–4839. 10.1073/pnas.1300436110.

Silverstone, A. L.; Ciampaglio, C. N.; Sun, T. The Arabidopsis RGA Gene Encodes a Transcriptional Regulator Repressing the Gibberellin Signal Transduction Pathway. The Plant Cell 1998, 10 (2), 155–169. 10.1105/tpc.10.2.155.

Silverstone, A. L.; Jung, H.-S.; Dill, A.; Kawaide, H.; Kamiya, Y.; Sun, T. Repressing a Repressor: Gibberellin-Induced Rapid Reduction of the RGA Protein in Arabidopsis. The Plant Cell 2001, 13 (7), 1555–1566. 10.1105/TPC.010047.

Spray, C. R.; Kobayashi, M.; Suzuki, Y.; Phinney, B. O.; Gaskin, P.; MacMillan, J. The Dwarf-1 (Dt) Mutant of Zea Mays Blocks Three Steps in the Gibberellin-Biosynthetic Pathway. Proc Natl Acad Sci U S A 1996, 93 (19), 10515–10518.

Sun, T. P.; Kamiya, Y. The Arabidopsis GA1 Locus Encodes the Cyclase Ent-Kaurene Synthetase A of Gibberellin Biosynthesis. Plant Cell 1994, 6 (10), 1509–1518.

Tal, Iris, Yi Zhang, Morten Egevang Jørgensen, Odelia Pisanty, Inês C. R. Barbosa, Melina Zourelidou, Thomas Regnault, et al. 2016. “The Arabidopsis NPF3 Protein Is a GA Transporter.” Nature Communications 7 (May): 11486. 10.1038/ncomms11486.

Tan, P.-H.; Zhang, L.; Yin, S.-X.; Teng, K. Heterologous Expression of a Novel Poa Pratensis Gibberellin 2-Oxidase Gene, PpGA2ox, Caused Dwarfism, Late Flowering, and Increased Chlorophyll Accumulation in Arabidopsis. Biologia plantarum 2018, 62 (3), 462–470. 10.1007/s10535-018-0788-1.

Thomas, S. G.; Phillips, A. L.; Hedden, P. Molecular Cloning and Functional Expression of Gibberellin 2- Oxidases, Multifunctional Enzymes Involved in Gibberellin Deactivation. Proceedings of the National Academy of Sciences 1999, 96 (8), 4698–4703. 10.1073/pnas.96.8.4698.

Ueguchi-Tanaka, M.; Ashikari, M.; Nakajima, M.; Itoh, H.; Katoh, E.; Kobayashi, M.; Chow, T.; Hsing, Y. C.; Kitano, H.; Yamaguchi, I.; Matsuoka, M. GIBBERELLIN INSENSITIVE DWARF1 Encodes a Soluble Receptor for Gibberellin. Nature 2005, 437 (7059), 693–698. 10.1038/nature04028.

Ueguchi-Tanaka, M., Y. Fujisawa, M. Kobayashi, M. Ashikari, Y. Iwasaki, H. Kitano, and M. Matsuoka.Rice Dwarf Mutant d1, Which Is Defective in the Alpha Subunit of the Heterotrimeric G Protein, Affects Gibberellin Signal Transduction. Proceedings of the National Academy of Sciences of the United States of America 2000 97 (21): 11638–43. 10.1073/pnas.97.21.11638.

Varbanova, M.; Yamaguchi, S.; Yang, Y.; McKelvey, K.; Hanada, A.; Borochov, R.; Yu, F.; Jikumaru, Y.; Ross, J.; Cortes, D.; Ma, C. J.; Noel, J. P.; Mander, L.; Shulaev, V.; Kamiya, Y.; Rodermel, S.; Weiss, D.; Pichersky, E. Methylation of Gibberellins by Arabidopsis GAMT1 and GAMT2. The Plant Cell 2007, 19 (1), 32–45. 10.1105/tpc.106.044602.

Vernoux, T.; Brunoud, G.; Farcot, E.; Morin, V.; Van den Daele, H.; Legrand, J.; Oliva, M.; Das, P.; Larrieu, A.; Wells, D.; Guédon, Y.; Armitage, L.; Picard, F.; Guyomarc’h, S.; Cellier, C.; Parry, G.; Koumproglou, R.; Doonan, J. H.; Estelle, M.; Godin, C.; Kepinski, S.; Bennett, M.; De Veylder, L.; Traas, J. The Auxin Signalling Network Translates Dynamic Input into Robust Patterning at the Shoot Apex. Mol Syst Biol 2011, 7, 508. 10.1038/msb.2011.39.

Wendt, Toni, Inger Holme, Christoph Dockter, Aileen Preuß, William Thomas, Arnis Druka, Robbie Waugh, Mats Hansson, and Ilka Braumann. HvDep1 Is a Positive Regulator of Culm Elongation and Grain Size in Barley and Impacts Yield in an Environment-Dependent Manner. PloS One 2016 11 (12): e0168924. 10.1371/journal.pone.0168924.

Wang, J.; Tian, C.; Zhang, C.; Shi, B.; Cao, X.; Zhang, T.-Q.; Zhao, Z.; Wang, J.-W.; Jiao, Y. Cytokinin Signaling Activates WUSCHEL Expression during Axillary Meristem Initiation. The Plant Cell 2017, 29 (6), 1373–1387. 10.1105/tpc.16.00579.

Wang, Q.; Kohlen, W.; Rossmann, S.; Vernoux, T.; Theres, K. Auxin Depletion from the Leaf Axil Conditions Competence for Axillary Meristem Formation in Arabidopsis and Tomato. The Plant Cell 2014, 26 (5), 2068–2079. 10.1105/tpc.114.123059.

Ward, J. M.; Smith, A. M.; Shah, P. K.; Galanti, S. E.; Yi, H.; Demianski, A. J.; van der Graaff, E.; Keller, B.; Neff, M. M. A New Role for the Arabidopsis AP2 Transcription Factor, LEAFY PETIOLE, in Gibberellin-Induced Germination Is Revealed by the Misexpression of a Homologous Gene, SOB2/DRN-LIKE. Plant Cell 2006, 18 (1), 29–39. 10.1105/tpc.105.036707.

Wild, M.; Davière, J.-M.; Regnault, T.; Sakvarelidze-Achard, L.; Carrera, E.; Lopez Diaz, I.; Cayrel, A.; Dubeaux, G.; Vert, G.; Achard, P. Tissue-Specific Regulation of Gibberellin Signaling Fine-Tunes Arabidopsis Iron-Deficiency Responses. Developmental Cell 2016, 37 (2), 190–200. 10.1016/j.devcel.2016.03.022.

Xie, Z.; Nolan, T. M.; Jiang, H.; Yin, Y. AP2/ERF Transcription Factor Regulatory Networks in Hormone and Abiotic Stress Responses in Arabidopsis. Frontiers in Plant Science 2019, 10.

Xu, Yanhao, Qiaojun Jia, Gaofeng Zhou, Xiao-Qi Zhang, Tefera Angessa, Sue Broughton, George Yan, Wenying Zhang, and Chengdao Li. Characterization of the sdw1 Semi-Dwarf Gene in Barley. BMC Plant Biology 2017, 17 (1): 11. 10.1186/s12870-016-0964-4.

Yaish, M. W.; El-kereamy, A.; Zhu, T.; Beatty, P. H.; Good, A. G.; Bi, Y.-M.; Rothstein, S. J. The APETALA-2-Like Transcription Factor OsAP2-39 Controls Key Interactions between Abscisic Acid and Gibberellin in Rice. PLOS Genetics 2010, 6 (9), e1001098. 10.1371/journal.pgen.1001098.

Yamaguchi, T.; Nagasawa, N.; Kawasaki, S.; Matsuoka, M.; Nagato, Y.; Hirano, H.-Y. The YABBY Gene DROOPING LEAF Regulates Carpel Specification and Midrib Development in Oryza Sativa. Plant Cell 2004, 16 (2), 500–509. 10.1105/tpc.018044.

Yu G (2022). enrichplot: Visualization of Functional Enrichment Result. R package version 1.16.1, https://yulab-smu.top/biomedical-knowledge-mining-book/.

Zhang, L.; Gu, L.; Ringler, P.; Smith, S.; Rushton, P. J.; Shen, Q. J. Three WRKY Transcription Factors Additively Repress Abscisic Acid and Gibberellin Signaling in Aleurone Cells. Plant Science 2015, 236, 214–222. 10.1016/j.plantsci.2015.04.014.

Zhang, X.; Wang, B.; Zhao, Y.; Zhang, J.; Li, Z. Auxin and GA Signaling Play Important Roles in the Maize Response to Phosphate Deficiency. Plant Science 2019, 283, 177–188. 10.1016/j.plantsci.2019.02.011.

Zhang, Y.; Zhang, B.; Yan, D.; Dong, W.; Yang, W.; Li, Q.; Zeng, L.; Wang, J.; Wang, L.; Hicks, L. M.; He, Z. Two Arabidopsis Cytochrome P450 Monooxygenases, CYP714A1 and CYP714A2, Function Redundantly in Plant Development through Gibberellin Deactivation. The Plant Journal 2011, 67 (2), 342–353. 10.1111/j.1365-313X.2011.04596.x.

Zhang, Z.; Zhang, J.; Chen, Y.; Li, R.; Wang, H.; Ding, L.; Wei, J. Isolation, Structural Analysis, and Expression Characteristics of the Maize (Zea Mays L.) Hexokinase Gene Family. Mol Biol Rep 2014, 41 (9), 6157–6166. 10.1007/s11033-014-3495-9.

Zhang, Z.-L.; Xie, Z.; Zou, X.; Casaretto, J.; Ho, T. D.; Shen, Q. J. A Rice WRKY Gene Encodes a Transcriptional Repressor of the Gibberellin Signaling Pathway in Aleurone Cells. Plant Physiol 2004, 134 (4), 1500–1513. 10.1104/pp.103.034967.

Zhao, X.; Yu, X.; Foo, E.; Symons, G. M.; Lopez, J.; Bendehakkalu, K. T.; Xiang, J.; Weller, J. L.; Liu, X.; Reid, J. B.; Lin, C. A Study of Gibberellin Homeostasis and Cryptochrome-Mediated Blue Light Inhibition of Hypocotyl Elongation. Plant Physiology 2007, 145 (1), 106–118. 10.1104/pp.107.099838.

Zhou, Z.; Sun, L.; Zhao, Y.; An, L.; Yan, A.; Meng, X.; Gan, Y. Zinc Finger Protein 6 (ZFP6) Regulates Trichome Initiation by Integrating Gibberellin and Cytokinin Signaling in Arabidopsis Thaliana. New Phytologist 2013, 198 (3), 699–708. 10.1111/nph.12211.

Zhu, Y.; Nomura, T.; Xu, Y.; Zhang, Y.; Peng, Y.; Mao, B.; Hanada, A.; Zhou, H.; Wang, R.; Li, P.; Zhu, X.; Mander, L. N.; Kamiya, Y.; Yamaguchi, S.; He, Z. ELONGATED UPPERMOST INTERNODE Encodes a Cytochrome P450 Monooxygenase That Epoxidizes Gibberellins in a Novel Deactivation Reaction in Rice. The Plant Cell 2006, 18 (2), 442–456. 10.1105/tpc.105.038455.

