## Supplemental Tables for "Comparing hormone dynamics in cereal crops via transient expression of hormone sensors"

**Table S1. GPS2 emission ratio in transfected leaf tissues in response to GA_3_ applications.**

| **GPS2 in leaf tissues** | | | | | | | | | | |
| --- | --- | --- | --- | --- | --- | --- | --- | --- | --- | --- |
|  | **Maize – B73 Rep #1** | | | | | **Maize – B73 Rep #2** | | | | |
| **[GA_3_] (uM)** | **0** | **0.1** | **1** | **10** | **100** | **0** | **0.1** | **1** | **10** | **100** |
| **Mean**  **±95%CI** | 2.19  ±0.10 | 2.06  ±0.05 | 2.54  ±0.09 | 3.07  ±0.10 | 3.16  ±0.09 | 1.81  ±0.10 | 1.34  ±0.10 | 2.35  ±0.09 | 2.79  ±0.16 | 2.17  ±0.06 |
| **Cliff’s Delta** | - | 0.00 | 0.22 | 0.50 | 0.54 | - | -0.52 | 0.39 | 0.63 | 0.32 |
| **N** | 405 | 1100 | 663 | 538 | 686 | 168 | 81 | 392 | 140 | 285 |
|  | **Maize – *d1* Rep #1** | | | | | **Maize – *d1* Rep #2** | | | | |
| **[GA_3_] (uM)** | **0** | **0.1** | **1** | **10** | **100** | **0** | **0.1** | **1** | **10** | **100** |
| **Mean**  **±95%CI** | 2.16  ±0.06 | **1.63**  ±0.15 | **1.53**  ±0.09 | **1.58**  ±0.05 | **-** | 1.83  ±0.15 | **1.47**  ±0.04 | **1.75**  ±0.18 | **1.42**  ±0.09 | **1.89**  ±0.11 |
| **Cliff’s Delta** | - | -0.45 | -0.58 | -0.56 | - | - | -0.35 | -0.06 | -0.46 | 0.07 |
| **N** | 578 | 80 | 185 | 583 | - | 96 | 635 | 64 | 178 | 166 |
|  | **Barley - Golden Promise Rep #1** | | | | | **Barley - Golden Promise Rep #2** | | | | |
| **[GA_3_] (uM)** | **0** | **0.1** | **1** | **10** | **100** | **0** | **0.1** | **1** | **10** | **100** |
| **Mean**  **±95%CI** | 1.32  ±0.04 | 2.14  ±0.13 | 2.15  ±0.13 | 2.12  ±0.15 | 2.78  ±0.13 | 1.20  ±0.03 | 1.50  ±0.05 | 1.87  ±0.13 | 1.83  ±0.09 | 2.22  ±0.08 |
| **Cliff’s Delta** | - | 0.72 | 0.60 | 0.64 | 0.86 | - | 0.46 | 0.53 | 0.72 | 0.88 |
| **N** | 219 | 202 | 211 | 157 | 323 | 447 | 349 | 247 | 208 | 291 |
|  | **Sorghum – BTx623 Rep #1** | | | | | **Sorghum – BTx623 Rep #2** | | | | |
| **[GA_3_] (uM)** | **0** | **0.1** | **1** | **10** | **100** | **0** | **0.1** | **1** | **10** | **100** |
| **Mean**  **±95%CI** | 2.09  ±0.13 | - | 3.38  ±0.14 | 3.16  ±0.38 | 3.19  ±0.29 | 2.59  ±0.10 | - | 2.92  ±0.31 | 2.93  ±0.18 | - |
| **Cliff’s Delta** | - | - | 0.68 | 0.50 | 0.71 | - | - | 0.21 | 0.14 | - |
| **N** | 119 | - | 242 | 54 | 41 | 188 | - | 34 | 156 | - |
|  | **Wheat – Kronos Rep #1** | | | | | **Wheat – Kronos Rep #2** | | | | |
| **[GA_3_] (uM)** | **0** | **0.1** | **1** | **10** | **100** | **0** | **0.1** | **1** | **10** | **100** |
| **Mean**  **±95%CI** | 1.15  ±0.06 | 1.61  ±0.31 | 1.45  ±0.52 | 1.81  ±0.98 | 1.38  ±1.18 | 1.24  ±0.07 | - | 2.30  ±0.18 | 1.42  ±0.12 | 2.21  ±0.11 |
| **Cliff’s Delta** | - | 0.29 | 0.15 | 0.21 | 0.40 | - | - | 0.79 | 0.18 | 0.82 |
| **N** | 101 | 42 | 12 | 7 | 4 | 86 | - | 69 | 75 | 168 |

**Table S2. GPS1-NR emission ratio in transfected leaf tissues in response to GA_3_ applications.**

| **GPS1-NR in leaf tissues** | | | | | | | | | | |
| --- | --- | --- | --- | --- | --- | --- | --- | --- | --- | --- |
|  | **Maize – B73 Rep #1** | | | | |  |  |  |  |  |
| **[GA_3_] (uM)** | **0** | **0.1** | **1** | **10** | **100** |  |  |  |  |  |
| **Mean**  **±95%CI** | 2.32  ±0.06 | 2.34  ±0.06 | 2.38  ±0.08 | 2.50  ±0.08 | 3.10  ±0.14 |  |  |  |  |  |
| **Cliff’s Delta** | - | 0.04 | 0.04 | 0.18 | 0.42 |  |  |  |  |  |
| **N** | 581 | 459 | 313 | 281 | 272 |  |  |  |  |  |
|  | **Maize – *d1* Rep #1** | | | | |  |  |  |  |  |
| **[GA_3_] (uM)** | **0** | **0.1** | **1** | **10** | **100** |  |  |  |  |  |
| **Mean**  **±95%CI** | 1.99  ±0.12 | 1.71  ±0.09 | 2.39  ±0.09 | 1.87  ±0.08 | - |  |  |  |  |  |
| **Cliff’s Delta** | - | -0.28 | 0.24 | -0.15 | - |  |  |  |  |  |
| **N** | 147 | 313 | 583 | 439 | - |  |  |  |  |  |
|  | **Barley - Golden Promise Rep #1** | | | | |  |  |  |  |  |
| **[GA_3_] (uM)** | **0** | **0.1** | **1** | **10** | **100** |  |  |  |  |  |
| **Mean**  **±95%CI** | 1.99  ±0.06 | 2.33  ±0.06 | 2.48  ±0.06 | 1.97  ±0.17 | 2.63  ±0.08 |  |  |  |  |  |
| **Cliff’s Delta** | - | 0.28 | 0.39 | -0.18 | 0.54 |  |  |  |  |  |
| **N** | 471 | 630 | 598 | 184 | 342 |  |  |  |  |  |
|  | **Sorghum – BTx623 Rep #1** | | | | | **Sorghum – BTx623 Rep #2** | | | | |
| **[GA_3_] (uM)** | **0** | **0.1** | **1** | **10** | **100** | **0** | **0.1** | **1** | **10** | **100** |
| **Mean**  **±95%CI** | 2.59  ±0.12 | - | 2.52  ±0.14 | 2.56  ±0.20 | 2.60  ±0.32 | 2.69  ±0.72 | - | 2.84  ±0.10 | 2.73  ±0.15 | - |
| **Cliff’s Delta** | - | - | -0.12 | -0.11 | 0.03 | - | - | 0.18 | 0.06 | - |
| **N** | 90 | - | 106 | 74 | 11 | 10 | - | 161 | 97 | - |
|  | **Wheat – Kronos Rep #1** | | | | | **Wheat – Kronos Rep #2** | | | | |
| **[GA_3_] (uM)** | **0** | **0.1** | **1** | **10** | **100** | **0** | **0.1** | **1** | **10** | **100** |
| **Mean**  **±95%CI** | 1.54  ±0.08 | 1.89  ±0.16 | 1.45  ±0.53 | 1.49  ±0.04 | 1.25  ±0.16 | 1.24  ±0.11 | - | 1.54  ±0.46 | 1.45  ±0.18 | 1.61  ±0.53 |
| **Cliff’s Delta** | - | 0.35 | -0.23 | -0.12 | -0.44 | - | - | 0.13 | 0.19 | 0.34 |
| **N** | 81 | 57 | 17 | 631 | 88 | 168 | - | 83 | 57 | 12 |

**Table S3. GPS2 emission ratio in transfected floral tissues in response to GA_3_ applications.**

| **GPS2 in floral tissues** | | | | | | | | | | |
| --- | --- | --- | --- | --- | --- | --- | --- | --- | --- | --- |
|  | **Maize – B73 Rep #1** | | | | |  |  |  |  |  |
| **[GA_3_] (uM)** | **0** | **0.1** | **1** | **10** | **100** |  |  |  |  |  |
| **Mean**  **±95%CI** | 1.80  ±0.13 | 1.79  ±0.11 | 1.95  ±0.17 | 2.48  ±0.20 | 3.00  ±0.21 |  |  |  |  |  |
| **Cliff’s Delta** | - | -0.03 | 0.05 | 0.32 | 0.52 |  |  |  |  |  |
| **N** | 188 | 272 | 192 | 196 | 223 |  |  |  |  |  |
|  | **Barley - Golden Promise Rep #1** | | | | | **Barley - Golden Promise Rep #2** | | | | |
| **[GA_3_] (uM)** | **0** | **0.1** | **1** | **10** | **100** | **0** | **0.1** | **1** | **10** | **100** |
| **Mean**  **±95%CI** | 1.67  ±0.05 | 2.12  ±0.07 | 2.82  ±0.17 | 3.20  ±0.08 | 2.64  ±0.10 | 1.51  ±0.05 | 1.89  ±0.08 | 2.70  ±0.09 | 2.81  ±0.09 | 2.67  ±0.13 |
| **Cliff’s Delta** | - | 0.38 | 0.63 | 0.85 | 0.61 | - | 0.46 | 0.53 | 0.72 | 0.88 |
| **N** | 348 | 439 | 263 | 586 | 451 | 319 | 254 | 396 | 538 | 167 |

| **GPS2 in floral tissues** | | | | | | | | | | |
| --- | --- | --- | --- | --- | --- | --- | --- | --- | --- | --- |
|  | **Maize – B73 Rep #1** | | | | |  |  |  |  |  |
| **[GA_3_] (uM)** | **0** | **0.1** | **1** | **10** | **100** |  |  |  |  |  |
| **Mean**  **±95%CI** | 1.80  ±0.13 | 1.79  ±0.11 | 1.95  ±0.17 | 2.48  ±0.20 | 3.00  ±0.21 |  |  |  |  |  |
| **Cliff’s Delta** | - | -0.03 | 0.05 | 0.32 | 0.52 |  |  |  |  |  |
| **N** | 188 | 272 | 192 | 196 | 223 |  |  |  |  |  |
|  | **Barley - Golden Promise Rep #1** | | | | | **Barley - Golden Promise Rep #2** | | | | |
| **[GA_3_] (uM)** | **0** | **0.1** | **1** | **10** | **100** | **0** | **0.1** | **1** | **10** | **100** |
| **Mean**  **±95%CI** | 1.67  ±0.05 | 2.12  ±0.07 | 2.82  ±0.17 | 3.20  ±0.08 | 2.64  ±0.10 | 1.51  ±0.05 | 1.89  ±0.08 | 2.70  ±0.09 | 2.81  ±0.09 | 2.67  ±0.13 |
| **Cliff’s Delta** | - | 0.38 | 0.63 | 0.85 | 0.61 | - | 0.46 | 0.53 | 0.72 | 0.88 |
| **N** | 348 | 439 | 263 | 586 | 451 | 319 | 254 | 396 | 538 | 167 |

**Table S4. GPS1-NR emission ratio in transfected floral tissues in response to GA_3_ applications.**

| **GPS1-NR in leaf tissues** | | | | | |
| --- | --- | --- | --- | --- | --- |
|  | **Maize – B73 Rep #1** | | | | |
| **[GA_3_] (uM)** | **0** | **0.1** | **1** | **10** | **100** |
| **Mean**  **±95%CI** | 2.21  ±0.13 | 2.46  ±0.15 | 2.44  ±0.22 | 2.32  ±0.28 | 2.65  ±0.34 |
| **Cliff’s Delta** | - | 0.22 | 0.11 | 0.04 | 0.08 |
| **N** | 149 | 119 | 122 | 64 | 93 |
|  | **Barley - Golden Promise Rep #1** | | | | |
| **[GA_3_] (uM)** | **0** | **0.1** | **1** | **10** | **100** |
| **Mean**  **±95%CI** | 2.47  ±0.16 | 2.23  ±0.09 | 2.68  ±0.20 | 2.50  ±0.19 | 2.96  ±0.09 |
| **Cliff’s Delta** | - | -0.21 | 0.19 | -0.03 | 0.33 |
| **N** | 85 | 282 | 57 | 142 | 529 |

**Table S5. GO-terms used in query for GA-related genes**

| **GO term** |  |
| --- | --- |
| GO:0009937 | regulation of gibberellic acid mediated signaling pathway |
| GO:0009938 | negative regulation of gibberellic acid mediated signaling pathway |
| GO:0009939 | positive regulation of gibberellic acid mediated signaling pathway |
| GO:0009739 | response to gibberellin |
| GO:0009740 | gibberellic acid mediated signaling pathway |
| GO:0042388 | gibberellic acid mediated signaling pathway, G-alpha-dependent |
| GO:0042390 | gibberellic acid mediated signaling pathway, G-alpha-independent |
| GO:0009728 | detection of gibberellic acid stimulus |
| GO:0071370 | cellular response to gibberellin stimulus |
| GO:0010476 | gibberellin mediated signaling pathway |
| GO:0009685 | gibberellin metabolic process |
| GO:0009686 | gibberellin biosynthetic process |
| GO:0033469 | gibberellin 12 metabolic process |
| GO:0045487 | gibberellin catabolic process |
