## Supplementary figures and images for "Comparing hormone dynamics in cereal crops via transient expression of hormone sensors"

### Supplemental Figure S1

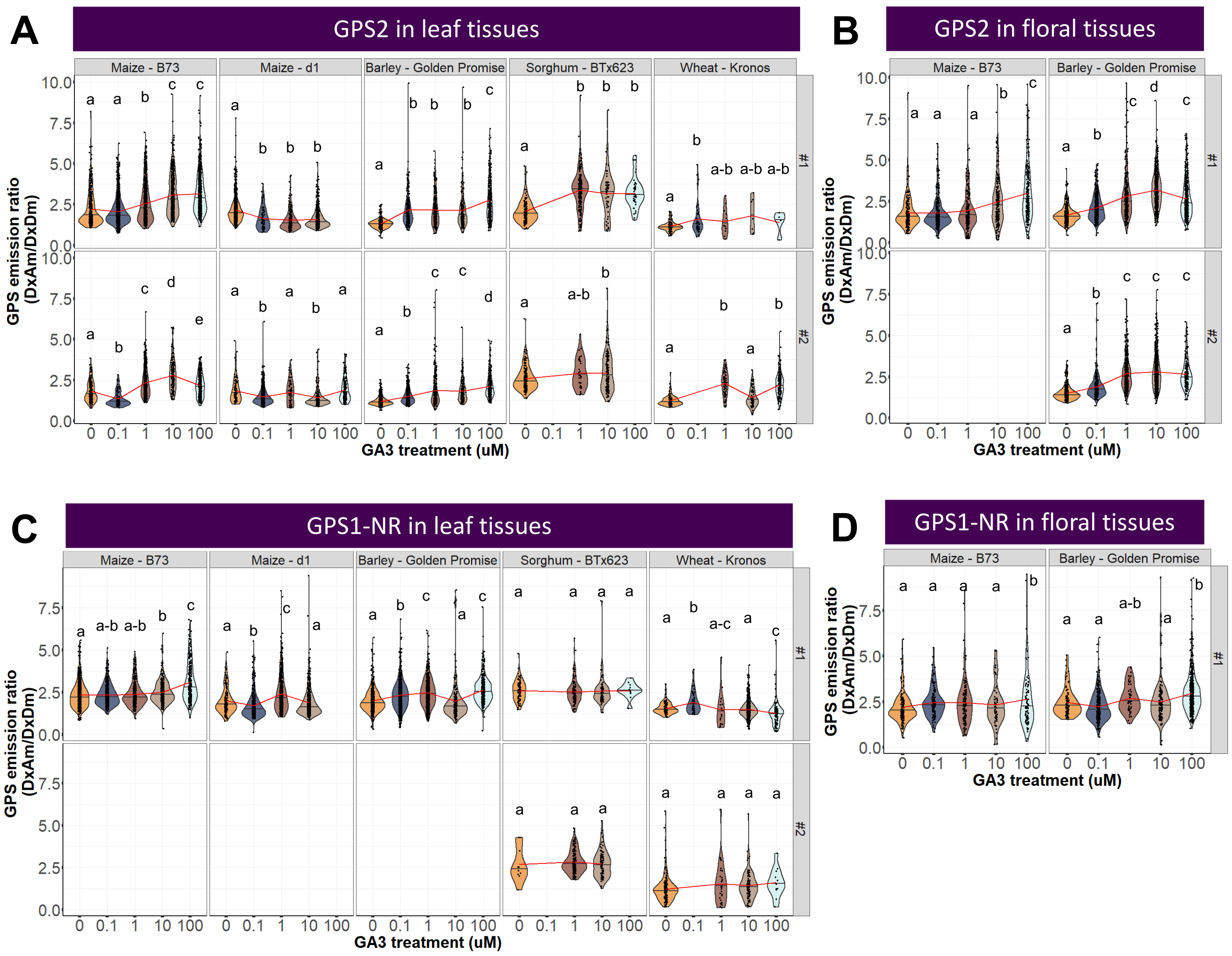
